# The dual character of the inhibitory functions of CD6

**DOI:** 10.1101/2022.04.29.490054

**Authors:** Rita F. Santos, Annika De Sousa Linhares, Marcos S. Cardoso, Ana Nóvoa, Hervé Luche, Fátima Gärtner, Bernard Malissen, Peter Steinberger, Simon. J. Davis, Moisés Mallo, Liliana Oliveira, Alexandre M. Carmo

## Abstract

T-cell membrane scaffold proteins play important roles in T cell biology, functioning as multi-functional signaling hubs. CD6 assembles a large intracellular signalosome but, unlike typical membrane-attached scaffolds like LAT or PAG, it has a sizeable ectodomain that binds a well-characterized ligand, CD166. It is unclear whether CD6 has net inhibitory or costimulatory functions or how its ectodomain influences these activities. To explore these questions, we dissected the signaling functions of the extracellular and cytoplasmic regions of CD6. We found that CD6 was delivered to the immunological synapse and suppressed T cell responsiveness *in vitro* wholly dependently of its cytoplasmic domain, indicating that CD6 very potently imposes tonic inhibition, acting as a structural and signaling inhibitory hub. However, the cell-intrinsic suppression of autoimmunity by CD6 *in vivo* was also impacted by extracellular interactions, demonstrated by the increased susceptibility of mice to experimental autoimmune encephalomyelitis after removal of the ligand binding region of the ectodomain of CD6. Our work identifies CD6 as a new class of ‘on/off switching’ scaffold-receptor that constrains immune responsiveness at two speeds. First, it sets signaling thresholds via tonic inhibition, functioning as a cytoplasmic membrane-bound scaffold and, second, by cycling between signaling-enabling and signaling-inhibiting ectodomain isoforms it functions as an immune checkpoint.

## INTRODUCTION

T cell receptor (TCR) recognition of antigen triggers the phosphorylation of immune receptor tyrosine activation motifs (ITAMs) in the CD3 complex by the SRC-family tyrosine kinases LCK and FYN (*1, 2*). This allows the SYK-family kinase ZAP70 to be recruited to the receptor (*3*), whereupon it phosphorylates a variety of downstream effectors, among the most important being proteins belonging to the transmembrane adaptor family. These scaffolding adaptors comprise a group of integral proteins that contain very small extracellular sections, including LAT, LAT2, PAG1 and TRAT1, among others, and they function more precisely as scaffolding anchors, assembling large intracellular regulatory complexes or “signalosomes” that launch multiple signaling cascades in response to immune receptor engagement (*4, 5*).

Scaffolds can generate either activating or inhibitory signaling outcomes that fine-tune the over-riding strength and nature of the stimulus. For example, LAT docks SH2 domain-containing enzymes and cytosolic adaptors such as PLCG1, PIK3R and GRB2, among others, promoting positive signaling (*6*). Conversely, PAG1 couples with CSK, the tyrosine kinase that phosphorylates inhibitory sites of LCK and FYN, therefore down-modulating the activity of these enzymes and signaling overall (*7*).

Another set of transmembrane proteins, exemplified by the structurally related CD5 and CD6 transmembrane receptors, have recently been considered to comprise an additional class of scaffolds insofar as they also assemble multi-component interactomes but have large extracellular domains (*8–11*). The signals that are relayed by CD5 repress T cell activation as it builds a signalosome composed mostly of inhibitory enzymes including CBL, CBLB, PTPN6 and RASA1 (*10, 12–14*). Thus far, it seems that CD5 signaling occurs independently of any extracellular interactions (*15*). Instead, the strength of CD5 inhibitory signaling is modulated by changes in its expression levels, increasing proportionally with the avidity to self-peptides during positive and negative selection (*16*). This may skew the peripheral TCR repertoire towards an effective recognition of foreign antigens as the exported thymic self-reactive clones that maintain high CD5 levels in the periphery produce optimal effector responses after pathogenic challenges (*17–20*).

In contrast to CD5, whether CD6 is co-stimulatory or inhibitory is controversial (*21–26*). CD6 has been shown to interact both with positive (*e.g*., LCK, ZAP70) (*9, 27*) and negative (*e.g*., PTPN6, UBASH3A) (*10, 11, 28*) regulators of T cell activation, as well as with many intracellular adaptors (*e.g*., LCP2, SH2D2A) (*29, 30*), but it has been considered historically to be a co-stimulatory molecule based on *in vitro* studies mostly using monoclonal antibodies (mAbs) targeting CD6 (*22–24, 31–34*). However, we have suggested that CD6 harbors substantial inhibitory activity based on experiments in which we manipulated the expression of CD6 in primary T cells and cell lines (*25*). The notion that CD6 is inhibitory is consistent with recent reports of CD6 knockout (CD6 KO) mice, in which purified CD6 KO T cells displayed enhanced responses following *in vitro* stimulation (*35, 36*). However, the *in vivo* effects of CD6 deficiency vary when tested in the context of different disease models. In the collagen-induced arthritis (CIA) model, CD6-null mice experienced more severe disease compared with wild-type (WT) mice (*36*), consistent with an activation-constraining role for CD6. On the contrary, in the experimental autoimmune encephalomyelitis (EAE) model of neuroinflammation, CD6 KO mice were completely refractory to the disease (*35*). Similarly, in experimental autoimmune uveitis (EAU) and in an imiquimod-induced model of psoriasis, the CD6-deficient animals exhibited significantly reduced disease compared with WT animals (*37, 38*). These disparities serve to highlight the complexity of the role CD6 plays in T cell physiology.

Also in contrast with CD5, CD6 signaling functions were assumed to be wholly dependent on the engagement of its ectodomain, positioning CD6 more as a linker, connecting extracellular-received stimuli to intracellular responses. The primary CD6 ligand is CD166 (ALCAM), which is widely expressed in many different tissues and cell types, including virtually all antigen presenting cells (APC) (*39, 40*). Upon the initial contact of T cells with APCs both CD6 and CD166 are redistributed to the immunological synapse (IS) (*22, 24*). Structurally similar to CD5, the extracellular region of CD6 comprises three scavenger receptor cysteine-rich (SRCR) domains (*41*), and it is through the SRCR membrane proximal domain, d3, that CD166 binding occurs (*42*). According to mRNA expression studies, the most abundant form of CD6 is encoded by all thirteen exons of the gene, but on T cell activation and also during thymic selection a CD6 isoform lacking d3 (CD6Δd3) is also expressed, and is therefore unable to bind to CD166 (*43*). The alternative splicing event that induces d3-encoding exon 5 skipping results from the impaired recruitment of the SRSF1 splicing factor, which would otherwise bind to intron 4 sequences promoting exon 5 inclusion (*44*). Importantly, a single nucleotide polymorphism (rs17828933) within the first intron of the *CD6* gene that is associated with higher expression of the CD6Δd3 isoform at the expense of the full-length isoform correlates with increased susceptibility to multiple sclerosis (MS) (*45, 46*).

Here, we address the function of CD6 by deleting the CD166-binding domain *in vivo* while leaving the signaling components untouched. We also examine *in vitro* the contribution of the extracellular and intracellular regions of CD6 to its IS redistribution, whole-cell adhesion and signal transduction. Our results show that CD6 is profoundly inhibitory and constrains immune responsiveness in two ways and at two levels, acting as both scaffold protein and immune checkpoint.

## RESULTS

### The impact of extra- and intra-cellular domains of CD6 on the regulation of T cell signaling

To determine the key components of CD6 that contribute to T cell signaling regulation, we constructed expression plasmids for the CD6WT molecule, the naturally-occurring isoform CD6Δd3, a cytoplasmic-truncation mutant (CD6Δcyt), and a binding- and signaling-disabled mutant that lacks both the d3 and the cytoplasmic tail (CD6Δd3Δcyt) (**Fig. 1A**). Je6-NF-κB::eGFP cells, a E6.1 Jurkat cell line engineered to express an eGFP reporter for the activation of the transcription factor NF-κB (*47*), were transduced to express these CD6 molecules at equivalent levels (**Fig. S1A**). Each of these cell lines was subjected to interaction either with Raji cells that naturally express CD166 (Raji-CD166^+^) or with Raji cells defective for CD166 expression (Raji-CD166^neg^) (*48*), in the absence or presence of superantigen (sAg).

**Fig. 1.**
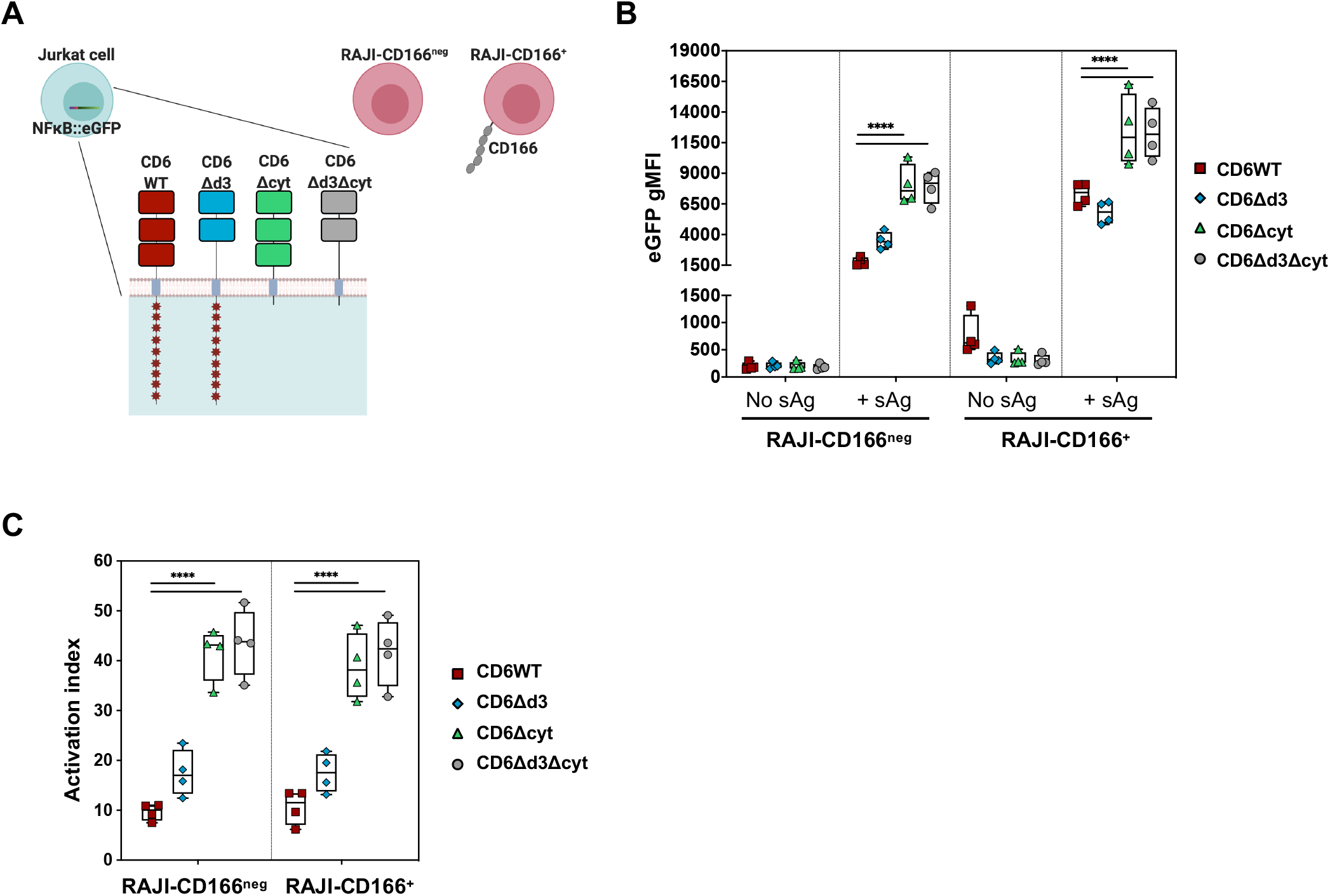
CD6 extra- and intra-cytoplasmic domains contribute differently to T cell activation. (**A**) Schematic representation of CD6 construc**t**s exp**r**essed in Je6-NF-κB::eGFP reporter cells. CD6 wild-type (WT), the naturally-occurring isoform lacking the ligand binding domain (CD6Δd3), a cytoplasmic-truncation mutant (CD6Δcyt), and a binding- and signaling-inert mutant (CD6Δd3Δcyt). Raji cells were used as antigen presenting cells, either naturally expressing CD166 (Raji-CD166^+^) or engineered to be defective of CD166 (Raji-CD166^neg^). (**B**) Je6-NF-κB::eGFP cells expressing CD6 WT and mutants were allowed to interact for 24 h at 37 °C with Raji-CD166^neg^ or Raji-CD166^+^ cells, previously incubated, or not, with the sAg staphylococcal enterotoxin E (SEE). T cell activation was assessed by flow cytometry analysis of NF-κB::eGFP up-regulation. (**C**) Normalized fold induction (activation index) of geometric mean of fluorescence intensity (gMFI) values of cells interacting in the presence of sAg over the gMFI values of the interactions without sAg. Each experiment was performed four times, with technical duplicates. Results of gMFI for each Je6-Raji pair condition in (B), and statistical analysis relative to the respective CD6WT in (C), ****, *p* < 0.001, two-way ANOVA, followed by Turkey’s multiple comparison test.

Je6 and Raji cells in different combinations were allowed to interact for 24 h, and the activation of NF-κB on the Je6 cells was assessed by measuring eGFP levels by flow cytometry. Activity profiles for each Je6:Raji pair are shown as raw data in **Fig. 1B**, and converted to activation index values (eGFP values of cells interacting in the presence of sAg / eGFP values of cells interacting in the absence of sAg) in **Fig. 1C**. The most conspicuous observation is that the cytoplasmic tail of CD6 strongly restrains the level of T cell activation, thus providing support to our previous studies (*25*). In particular, cells that express CD6 mutants lacking the cytoplasmic tail (CD6Δcyt and CD6Δd3Δcyt) were significantly more activated than those expressing tail-containing CD6 (CD6WT and CD6Δd3) (**Fig. 1C**).

Rather surprisingly, however, the interaction of the extracellular domain of CD6 with Raji-expressed CD166 did not induce a step-change in the activation, or inhibition, of the Je6 cells. As anticipated, and given that they do not express the ligand-binding domain, Je6-NF-κB::eGFP-CD6Δd3 cells showed the same activation whether interacting with CD166-negative or CD166-positive Raji cells (**Fig. 1C**). However, the same behavior was unexpectedly observed for the Je6-NF-κB::eGFP-CD6WT cells, *i.e*., CD6WT-expressing Je6 cells displayed exactly the same activation index regardless of interacting with CD166-defficient or with CD166-expressing Raji cells (**Fig. 1C**).

Overall, using a cellular system where model Jurkat T cells are allowed to directly interact with APCs without the interference of activating or blocking mAbs or other reagents, we can conclude that upon activation triggered by sAg presentation the cytoplasmic tail of CD6 is a major contributor for the inhibitory effects and that binding of CD6 to CD166 expressed on APCs has no major impact on signaling strength.

### The role of the 3^rd^ SRCR domain of CD6 on cell adhesiveness, immunological synapse organization and signaling inhibition

Although it appeared that the physical binding of CD6WT to CD166 led to no differences on activation when compared with the absence of that interaction using CD6Δd3, we noticed that in the absence of sAg (*i.e*., non-activated), the eGFP levels of only the Je6-CD6WT cells interacting with Raji-CD166^+^ cells were higher from the start (~ 2.3-fold), when compared with all other CD6-mutant-expressing cells (**Fig. 1B**, plots in Raji-CD166^+^/no sAg). This effect suggested that binding of CD6 to CD166 might actually have an impact not during activation but, instead, prior to antigen presentation. We thus addressed the relevance of the CD6-CD166 pair on cell adhesiveness during antigen-dependent and -independent priming of T cells.

E6.1 Jurkat cells expressing CD6WT were placed in contact with either Raji-CD166^neg^ or Raji-CD166^+^ cells in the absence or presence of sAg, and the percentage of cell conjugates (doublets) was measured by imaging flow cytometry (representative images shown in **Fig. 2A**, left). Whereas following sAg presentation (activated cells) the presence of CD166 resulted in no changes in the number of doublets, in the case when no sAg was present the percentage of E6.1-CD6WT cells that conjugated with Raji-CD166^+^ cells was significantly higher than those conjugating with Raji-CD166^neg^ cells (**Fig. 2A**, right). When using E6.1 cells expressing CD6Δd3, this effect was not reproduced (**Fig. 2B**). These results confirm that the contribution of the CD6-CD166 ligation towards cell adhesion is mostly relevant still in the absence of TCR engagement.

**Figure 2.**
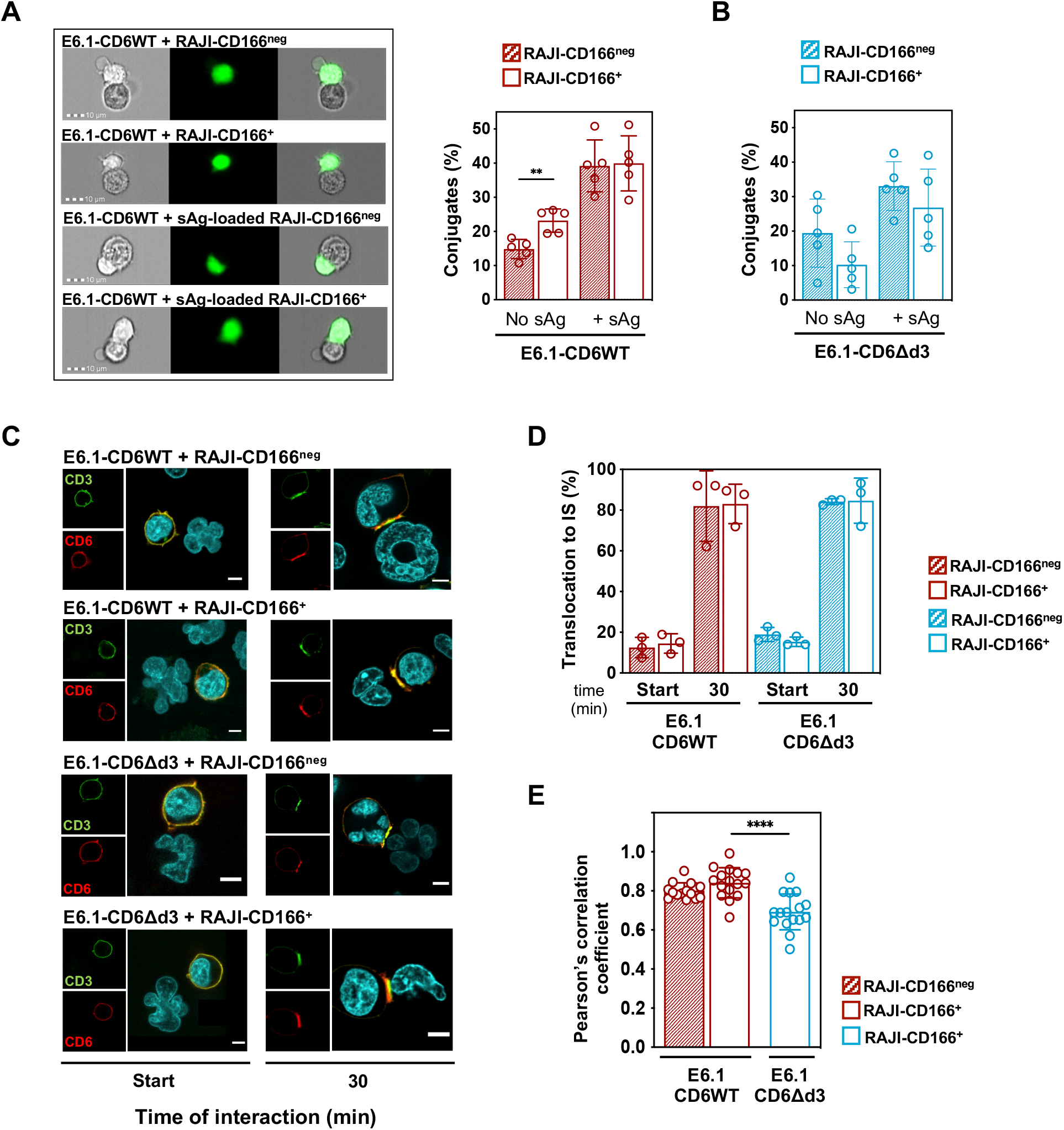
CD6 translocation to the IS is independent of binding to CD166, but the CD6-CD166 interaction impacts on antigenindependent T cell:APC adhesion. (**A**) E6.1 Jurkat CD6WT cells, previously stained with a cell tracker dye (CMFDA), were conjugated with Raji-CD166^neg^ or with Raji-CD166^+^ cells, pre-loaded or not with sAg. Left, representative images of cell:cell fixed conjugates in the absence or presence of CD6-CD166 engagement, assessed by imaging flow cytometry. Right, graph shows the percentage of conjugates in all stained cells from five independent experiments with the representative mean ± SD. **, *p* < 0.01, unpaired Student’s t test with Welch’s correction. (**B**) Percentage of conjugates formed between E6.1-CD6Δd3 and Raji-CD166^neg^ or Raji-CD166^+^ cells, preloaded or not with sAg, from five independent experiments. (**C**) Representative confocal images of CD6WT and CD6Δd3 localization at immunological synapses formed between E6.1 and Raji-CD166^neg^ or Raji-CD166^+^ cells. Cells were fixed within the first 5 min of initiation of contacts (left images) and after 30 min of cell interactions (right images). CD3 is shown in green and defines the cSMAC, CD6 (detected by MEM98, anti-CD6d1) is visualized in red. Merge images allow to detefine CD3 and CD6 co-localization. Scale bar, 5 μm. (**D**) Frequency of conjugates in which CD6WT and CD6Δd3 accumulate at the contact zone. Results are from at least 40 conjugates per condition, obtained from at least three independent experiments. Quantification was done by blind analysis from three different people. (**E**) Quantitative analysis of CD3 and CD6 pixel co-localization upon conjugate formation between sAg-loaded Raji and E6.1 cells expressing CD6WT or CD6Δd3. Pearson’s correlation coefficients from the synaptic region were generated using ImageJ software and JACoP plugin. Values obtained from conjugates from three independent experiments are represented as mean ± SD. Representative 3D projections are shown in supplementary information (Movie S1). ****, *p* < 0.001, unpaired Student’s t test with Welch’s correction.

One other unforeseen observation from the data on Fig. 1 was that although both natural isoforms, CD6WT and CD6Δd3, contain the same intracellular signaling motifs, Je6-CD6WT cells were slightly more repressed than Je6-CD6Δd3 cells even when interacting with Raji-CD166^neg^ cells (*p* = 0.029, Mann-Whitney test for direct comparisons), a condition in which none of the two CD6 isoforms is able to contact the ligand. We therefore investigated whether these isoforms had different distributions in the immunological synapse (IS) that could explain their different activation profiles.

E6.1 Jurkat cells expressing CD6WT or CD6Δd3 were put in contact with sAg-loaded Raji-CD166^neg^ or Raji-CD166^+^ cells, and CD6 molecules were visualized by immunofluorescence. Within the first 5 min of cell contacts, CD6WT and CD6Δd3, as well as CD3, were evenly distributed at the whole cell surface in all conditions [**Fig. 2C**, left panels (Start)], although in a small percentage of cells an enrichment of CD6WT or CD6Δd3 could already be detected at the E6.1:Raji interfaces (**Fig. 2D**). After 30 min of cell contacts, in E6.1-CD6WT cells interacting with Raji-CD166^+^ cells CD6 was transported to the interface between the two cells, as expected [**Fig. 2C**, 2^nd^ row, right panels (30 min); **Fig. 2D**, red-bordered column]. Unexpectedly, CD6WT also targeted to the IS of E6.1-CD6WT cells interacting with Raji-CD166^neg^ cells (**Fig. 2C**, 1^st^ row, 30 min; **Fig. 2D**, red column), indicating that the synaptic localization of CD6 occurs independently of binding to CD166, in complete contrast with the current paradigm (*24, 43*). This conclusion was further strengthened by the observation that in Jurkat cells conjugated with either Raji-CD166^neg^ or Raji-CD166^+^ cells, CD6Δd3 was also directed to the IS (**Fig. 2C**, 3^rd^ and 4^th^ rows, 30 min; **Fig. 2D**, blue and blue-bordered columns).

Quantitative analysis of CD6 co-localization with CD3 revealed that, however, the precise localization and interaction with the TCR signaling machinery is not identical between CD6WT and CD6Δd3 (**Fig. 2E** and **Movie S1**). In particular, CD6Δd3 co-localization with CD3 was significantly lower than that of CD6WT. On the other hand, CD6WT showed similar levels of interaction or proximity with CD3 independently of the presence of CD166 in the APC. These results suggest that the SRCR domain 3 of CD6 may establish *cis*-interactions with the TCR complex independently of binding CD166 on an opposing cell, being this way an inhibitory mediator that, by interacting directly or indirectly with the TCR signaling apparatus, helps to restrain T cell activation.

### Translocation of CD6 to the immunological synapse is mediated by the cytoplasmic tail, which additionally contains multiple inhibitory signaling motifs

Given that the extracellular domain was not mandatory for the synaptic localization of CD6, we questioned whether the translocation to the IS could depend on the cytoplasmic tail. E6.1 cells expressing CD6Δcyt were allowed to contact sAg-loaded Raji-CD166^neg^ or Raji-CD166^+^ cells, and the localization of CD6Δcyt was visualized by immunofluorescence. CD6Δcyt was dispersed throughout the cell surface and only concentrated at the IS in a very small fraction of cell contacts within the first 5 min. However, unlike CD6WT or CD6Δd3, the bulk of CD6Δcyt did not translocate to mature synapses (defined by CD3 enrichment after 30 min of cell contacts) even though it contains the ligand-binding domain (**Fig. 3A, B**). Nonetheless, we assessed whether CD6Δcyt could play a role in antigen-independent or -dependent adhesion of Jurkat cells. E6.1 Jurkat cells expressing CD6Δcyt were placed in contact with Raji-CD166^neg^ or Raji-CD166^+^ cells in the absence or presence of sAg, and the percentage of cell conjugates was measured by imaging flow cytometry. As can be seen in **Fig. 3C**, CD6Δcyt did not induce significant differences in the number of doublets formed between E6.1-CD6Δcyt and Raji-CD166^neg^ or Raji-CD166^+^ cells, either in the absence (left columns) or the presence (right columns) of sAg. These observations confirm that the IS-targeting motifs are contained within the cytoplasmic tail of CD6, and lack of the tail impairs both the CD6 translocation to the IS and any major contribution to cellular adhesion.

**Fig. 3.**
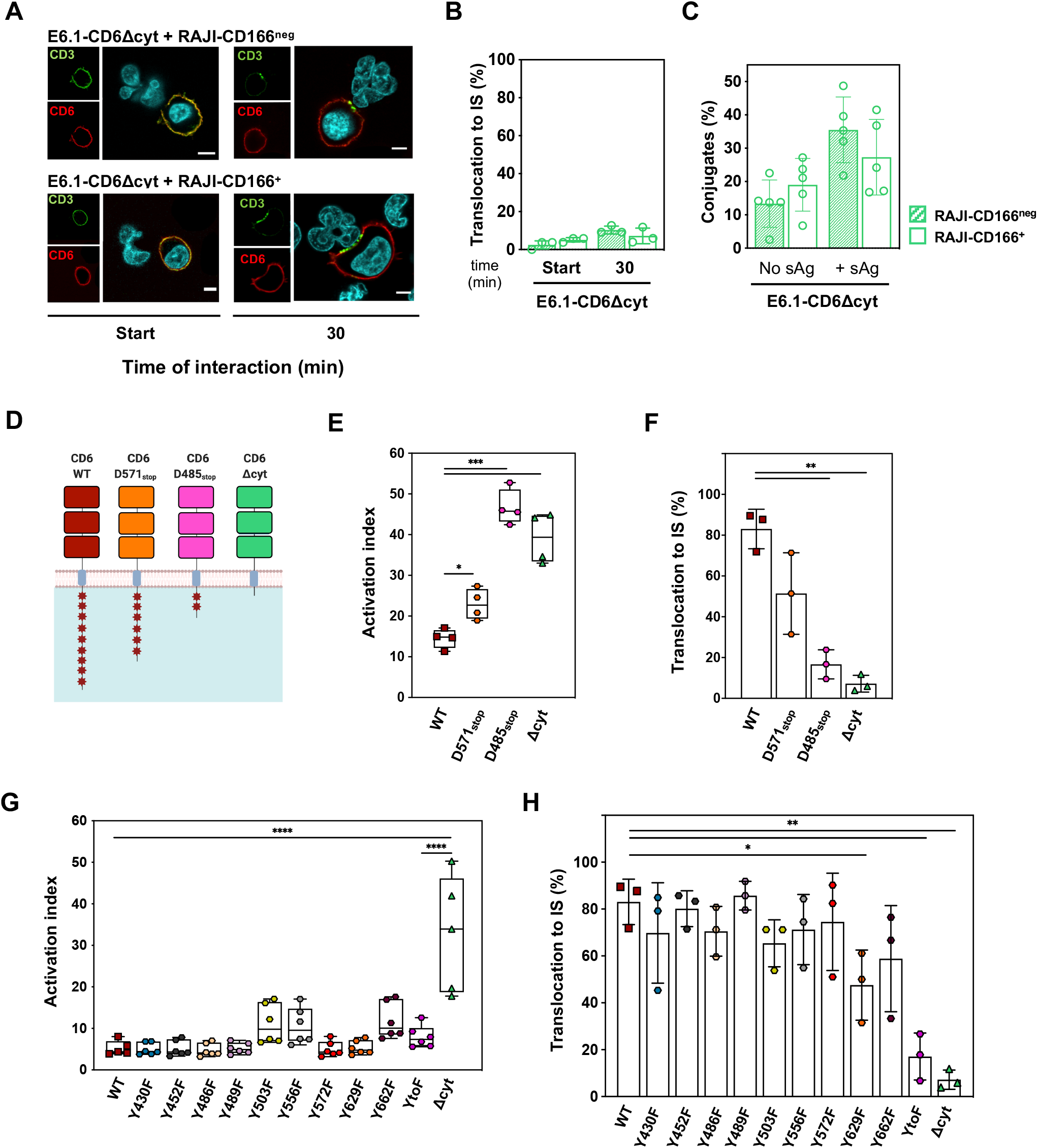
CD6 translocation to the IS and signaling inhibition are dependent on the cytoplasmic tail. (**A**) Representative confocal images of conjugates formed between sAg-loaded Raji-CD166^neg^ or Raji-CD166^+^ cells with E6.1 cells expressing CD6Δcyt. Cells were fixed within the first 5 min of initiation of contacts (left images) and after 30 min of cell interactions (right images). Immunostainings of CD3 and CD6 (using MEM98) were performed, CD3 is shown in green and defines the cSMAC, CD6 is displayed in red. Merge images reveal no specific co-localization between CD3 and CD6Δcyt. Scale bar: 5 μm. (**B**) Frequency of conjugates in which CD6Δcyt accumulates at the contact zone. Differences not significant, oneway ANOVA, followed by Dunnett’s multiple comparison test and unpaired Student’s t test with Welch’s correction. (**C**) Percentage of conjugates, measured by imaging flow cytometry, formed between E6.1-CD6Δd3 and Raji-CD166^neg^ or Raji-CD166^+^ cells, pre-loaded or not with sAg, from five independent experiments, with the representative mean ± SD. Differences not significant, unpaired Student’s t test with Welch’s correction. (**D**) Schematic representation of CD6 constructs expressed in Je6-NF-κB::eGFP reporter cells, highlighting the number of tyrosine residues (red asterisks) remaining in each CD6-truncation mutant. (**E**) Je6-NF-κB::eGFP cells expressing CD6 cytoplasmic tail truncations CD6D571_stop_, CD6D485_stop_ and CD6Δcyt were co-cultivated with Raji-CD166^+^ cells, previously incubated without or with sAg. Activation indexes are represented in the graph. (**F**) Frequency of synapses in which CD6 accumulates at the contact zone formed between sAg-loaded Raji and E6.1 cells expressing the cytoplasmic tail truncations. Cells were fixed after 30 min of interaction. (**G**) Je6-NF-κB::eGFP cells expressing different CD6 mutants, each having a Y-to-F substitution, or a mutant with all nine CD6 tyrosine residues substituted by phenylalanines (YtoF) were co-cultivated with Raji-CD166^+^ cells, previously incubated without or with sAg. Activation indexes are represented in the graph. (**H**) Percentage of synapses in which CD6 accumulates at the contact zone between Raji-CD166^+^ cells loaded with sAg and E6.1 cells expressing CD6 with tyrosine to phenylalanine substitutions. (E,G) Each experiment was performed four times, with technical duplicates. Statistical differences shown only for CD6WT. *, *p* < 0.05; ****, *p* < 0.001, two-way ANOVA, followed by Turkey’s multiple comparison test. (B,F,H) Results are from at least 40 conjugates per condition, obtained from at least three independent experiments. Quantification was done by blind analysis from three different people. *, *p* < 0.05; **, *p* < 0.01, one-way ANOVA, followed by Dunnett’s multiple comparison test and unpaired Student’s t test with Welch’s correction.

To further dissect the inhibitory and organizational components of the CD6 cytoplasmic domain we constructed mutants with progressively reduced tail length (**Fig. 3D**). Je6-NF-κB::eGFP cells expressing CD6WT or the cytoplasmic truncation mutants CD6-D571_stop_, CD6-D485_stop_ or CD6Δcyt were allowed to interact for 24 h with Raji-CD166^+^ cells in the absence or presence of sAg, and the equivalent E6.1 cells expressing the same mutants allowed to conjugate for 30 min with sAg-loaded Raji-CD166^+^ cells. The eGFP levels were measured for each condition, the activation indexes calculated, and the percentage of conjugates in which CD6 translocated to the IS was assessed. Overall, there was a correlative tendency between shorter CD6 cytoplasmic tail and lower sAg-independent priming of Jurkat cells [**Fig. S2A**, left (No sAg)], less repressed cell responses (**Fig. 3E**) and fewer cells with CD6 localized at the IS (**Fig. 3F**). These results show that both the inhibitory motifs of CD6 as well as those that determine its targeting to the IS are spread along the cytoplasmic tail. Moreover, they prove that, given that all mutants contain the CD166-binding domain, the less CD6 is localized at the synapse the lower is the adhesive impact of the molecule on the sAg-independent cell priming. Nevertheless, these data also raised the possibility that the same motifs were responsible for the conjugation of both effects and eventually that the inhibitory role of CD6 could be highly dependent on its localization at the T:APC interface and not solely determined by interactions with inhibitory effectors.

The cytoplasmic domain of CD6 contains nine tyrosine residues that upon phosphorylation may mediate interactions with SH2 domain- or phosphotyrosine binding (PTB) domain-containing enzymes or adaptors that participate in signal transduction and/or help in assembling a multi-protein complex. We constructed CD6 mutants with individual Y-to-F substitutions, and analyzed both their signaling properties and their capacity to target CD6 to the IS. Most of the CD6 Y-to-F mutations had no detectable impact on signal transduction, as measured by eGFP levels in Je6-NF-κB::eGFP cells when put into contact with Raji-CD166^+^ cells in the absence or presence of sAg (**Fig. 3G** and **Fig. S2B**). However, substitutions of three tyrosine residues, Y503, Y556 and Y662, resulted in a slight increase in the activation index, *i.e*., losses of inhibitory signaling, indicating that these specific tyrosines likely play a role in the inhibitory effect on signal transduction. The simultaneous substitution of all 9 tyrosine residues resulted, however, in a CD6 molecule that only residually differed in their inhibitory properties from the WT molecule. This may suggest that stimulatory effectors can conceivably bind to some of the phosphorylated tyrosine residues; disruption of their binding could result in the abrogation of any potential co-stimulatory signaling, rendering on aggregate of all CD6 tyrosine substitutions a close to neutral effect. By contrast, the CD6Δcyt mutant, which also lacks all 9 tyrosine residues, was much less repressed than the CD6-YtoF molecule. This shows that the cytoplasmic tail must contain phosphotyrosine-independent inhibitory motifs that very strongly repress activation.

Translocation of CD6 Y-to-F mutants to the IS was also measured in E6.1 cells, co-cultured with Raji-CD166^+^ cells pre-incubated with sAg (**Fig. 3H**). Overall, the different mutants showed only slight variations in the IS-localization when compared with that of CD6WT, with the exception of the Y629F substitution that reduced IS-localization by approximately 40%. Interestingly, when all tyrosine residues were replaced by phenylalanines the decline in IS-targeting was very pronounced. This suggests that extensive tyrosine phosphorylation of the CD6 cytoplasmic tail is essential to build the CD6 interactome that brings this molecule to the IS, and no specific tyrosine substitution is able to singlehandedly deconstruct the complex. Given that no single point mutation had major effects on both signaling inhibition and synaptic translocation, we can conclude that these are two independent phenomena probably mediated by non-coincident motifs of the cytoplasmic tail. Importantly, because CD6-YtoF is still a strong inhibitor but does not target to the IS, it becomes evident that CD6 is not required to localize at the synapse to exert its inhibitory function.

Overall, these data show that the cytoplasmic domain of CD6 contains non-overlapping phosphorylation-dependent sequences that individually are moderately relevant. The whole of the cytoplasmic tail is, nonetheless, absolutely crucial to the targeting of CD6 to the IS and in mediating strong inhibitory signaling.

### Effects of the omission of the 3^rd^ SRCR domain of CD6 on thymopoiesis

The two physiological CD6 isoforms that are known to mostly impact on CD6 function are the predominant full-length WT form, and CD6Δd3 which is enriched in activated T cells and developing thymocytes (*43*). However, *in vivo* models of CD6 may be particularly hard to interpret given that, paradoxically, it is the endogenous molecule in WT mice that undergoes cyclical modifications, and currently there is no technology that can determine in a WT mouse when and where during thymocyte development or in the course of an immune response CD6 is being expressed as a full-length protein or as the CD6Δd3 isoform. The molecular framework that we developed using the Jurkat cell model might, therefore, be of particular interest in helping to interpret the events observed in the *in vivo* experiments.

We addressed the biological relevance of d3 by developing the CD6Δd3 mouse model, thus creating a near-physiological environment where, while abrogating d3-dependent ligations, namely in *trans* with APC-expressed CD166 and in *cis* with the TCR complex, all other inherent functions of CD6 can operate in their natural setting. CD6Δd3 C57BL/6 mice were generated by deleting *CD6* exon 5 through CRISPR/Cas9 engineering (**Fig. 4A**). The deletion of d3 was confirmed by PCR using primers spanning the target sites (**Fig. 4B**) and by western blotting using a mouse CD6d1 mAb, which confirmed the decreased CD6 protein size in the mutant mice (**Fig. 4C**).

**Fig. 4.**
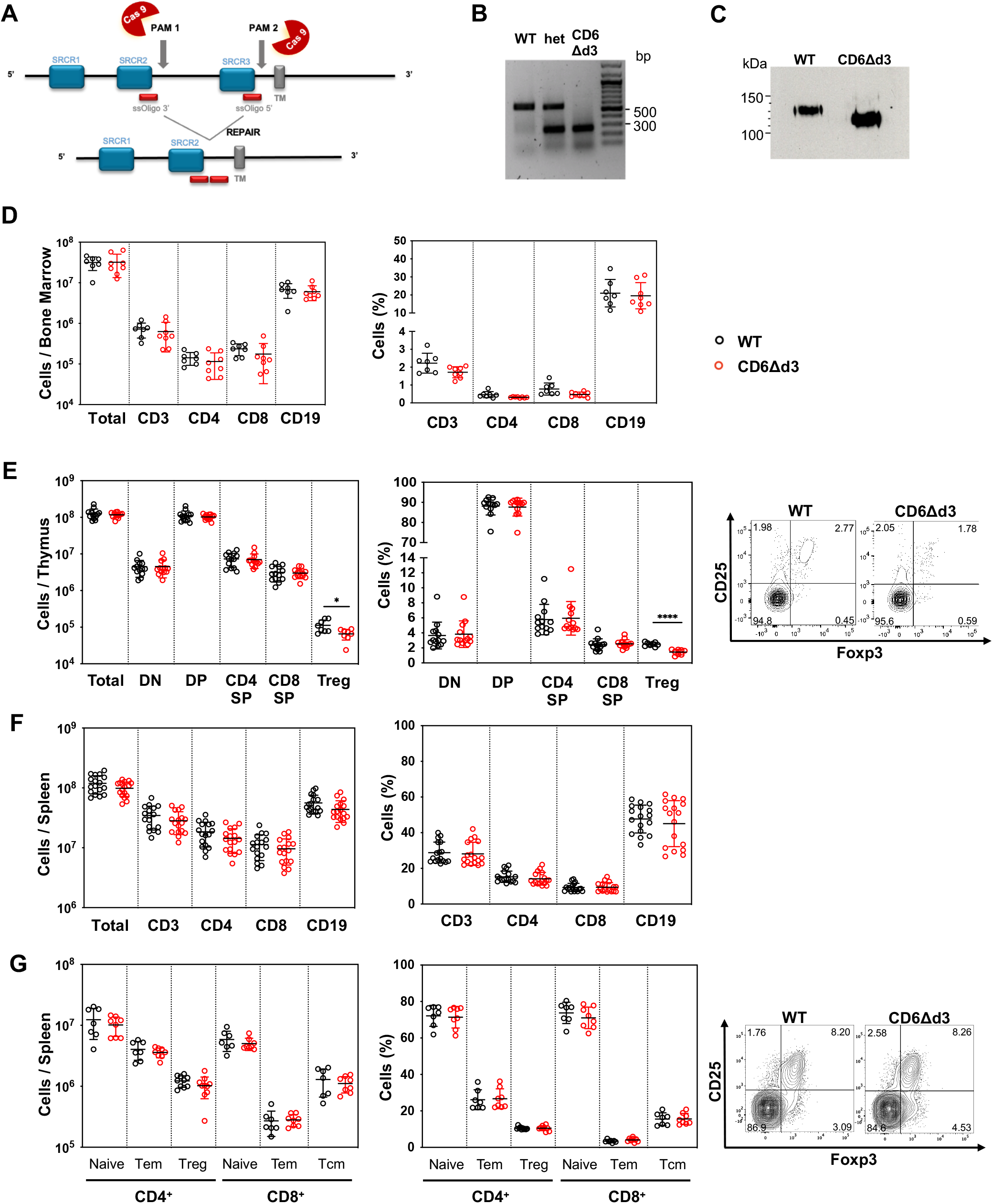
Development and immunophenotyping characterization of the C57BL/6 CD6Δd3 mouse. (**A**) Schematic representation of the CRISPR/Cas9 *Cd6* genome editing strategy to exclude the SRCR domain 3 of CD6, encoded by exon 5, by targeting two PAM sequences in the adjacent introns 4 and 5. (**B**) PCR using forward and reverse primers oligomerizing to sequences in the exon 4/intron 4 boundary, and within intron 5, respectively (the difference in size comparing inclusion *vs*. exclusion is 291 bp). (**C**) WB using an anti-CD6d1 mAb confirming the deletion of d3 in the CD6Δd3 mouse. (**D-G**) Flow cytometry analysis of total cell numbers (left) and frequencies (right) in WT (black circles) and CD6Δd3 (red circles) mice of: (**D**) bone marrow lymphoid cell populations; (**E**) thymocyte subsets (DN, double negative; DP, double positive; SP, single positive CD4^+^ or CD8^+^; Treg, T regulatory). In the far right, representative contour plot of Foxp3^+^CD25^+^ thymocytes amongst the CD4^+^ SP total; (**F**) spleen lymphoid cell populations; and (**G**) CD4^+^ and CD8^+^ T cell subsets in the spleen [naïve: CD62L^high^CD44^low^; Tem (T effector memory): CD62L^low^CD44^high^; Treg: Foxp3^+^CD25^+^; Tcm (T central memory): CD62L^high^CD44^high^]. In the far right, representative contour plot of Foxp3^+^CD25^+^ CD4^+^ splenocytes. Each dot represents one individual mouse, and horizontal lines indicate the respective mean ± SD from at least three independent experiments. *, *p* < 0.05; ****, *p* < 0.001, unpaired Student’s t test with Welch’s correction.

In depth immune-phenotyping analyses performed in spleen and thymus of 12-week-old WT and CD6Δd3 mice by multi-parametric flow cytometry showed no major differences in leukocyte populations, except a slight increase in the proportion of macrophages in the spleen and a lower frequency of CD69^+^ thymocytes among the CD3^-^ double positive (DP) population in CD6Δd3 mice, compared with control mice (**Fig. S3**). Absolute numbers and frequencies of individual lymphoid cell populations in the bone marrow, thymus and spleen showed no additional differences (**Fig. 4D**), namely of different subsets of mature/effector T cells, between sex- and age-matched CD6WT and CD6Δd3 littermates, except for a noticeable reduction in the frequency and also of total numbers of CD4^+^CD25^+^Foxp3^+^ T regulatory (Treg) cells in the thymus of CD6Δd3 mice (**Fig. 4E**), though not in the spleen (**Fig. 4G**).

Given that in steady state adult mice the only difference we detected in T cell development was the lower abundance of Tregs in thymi of CD6Δd3 mice, we performed a phenotypic characterization of the effect of lack of d3 on T cell maturation in much younger mice (one-week-old), in which thymic function is highly active. The numbers of the different thymocyte developmental subsets were imbalanced in young CD6Δd3 mice compared with young WT mice (**Fig. 5A**), in contrast to what we had observed in the adults. The decreased number of CD4^+^ and CD8^+^ single-positive (SP) cells in CD6Δd3 mice suggests a deficiency in thymocyte selection, leading to a reduction in the number of cells reaching the end of maturation in the mutant mice. The generation of Tregs, a process that occurs concomitantly with negative selection, was also affected, with a pronounced decrease in the number of Tregs in CD6Δd3 young mice (**Fig. 5A**). In addition, we observed a large decrease in the frequency and numbers of CD3^+^CD69^-^ and CD3^+^CD69^+^ cells in the thymi of CD6Δd3 young mice, markers that are correlated with late stages of maturation and selection (**Fig. 5B**).

**Fig. 5.**
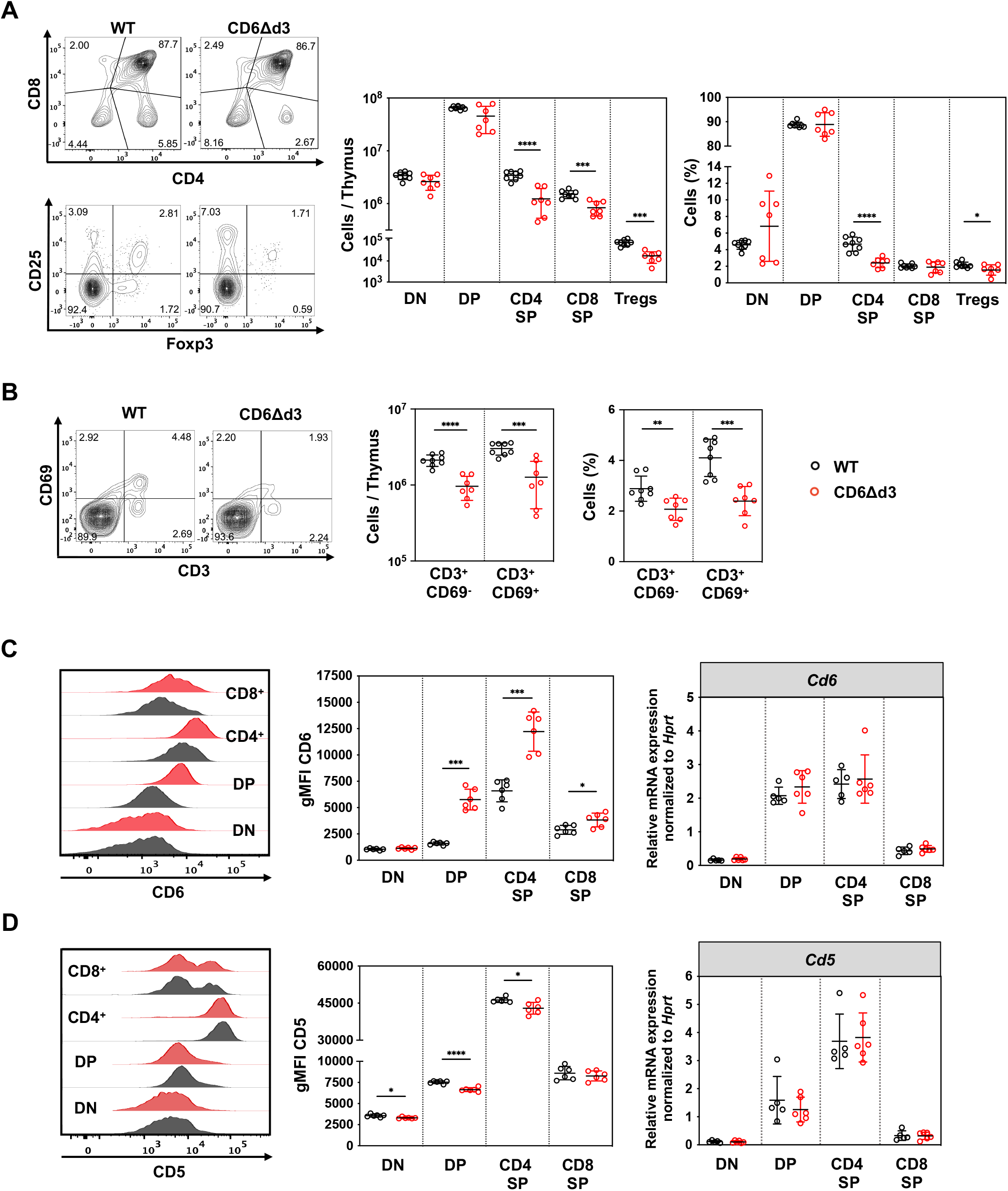
Impact of the absence of CD6d3 on T cell maturation in thymuses of one-week-old mice. (**A**) Total number and frequencies of cells in each thymocyte subset from WT (black circles) and CD6Δd3 (red circles) thymuses. Representative contour plots on the left (DN, CD8^+^ SP, DP and CD4^+^ SP T cells, clockwise in the top panels; Tregs, top-right quadrant in bottom panels). The corresponding graphs are shown on the right. (**B**) On the left, representative contour plots showing the percentage of cells that rearranged a pre-TCR receptor (CD3^+^) and that were positively selected (CD3^+^CD69^+^) amongst total live thymocytes. The corresponding graphs are shown on the right. (**C-D**) Expression of CD6 (C) and CD5 (D) in the different thymocyte subsets of WT and CD6Δd3 mice. On the left, representative histograms of CD6 surface expression (staining with mAb BX222 for CD6d1) or CD5 surface expression, measured by flow cytometry in DN, DP, CD4^+^ SP and CD8^+^ SP T cell populations, with the correspondent graphs in the middle panels indicating gMFI of CD6 and CD5 for each individual mouse. On the right panels, it is indicated the relative mRNA expression of *Cd6* and *Cd5* in sorted thymic cells of one-week-old mice normalized to *Hprt* mRNA expression. Each dot represents one individual mouse, and horizontal lines indicate the respective mean ± SD. *, *p* < 0.05; **, *p* < 0.01; ***, *p* < 0.005; ****, *p* < 0.001 (unpaired Student’s t test with Welch’s correction). Each experiment was performed at least twice. DN: double negative; DP: double positive; SP: single positive; gMFI: geometric mean fluorescence intensity.

CD6 levels increase concomitantly with T cell maturation (*49*). We observed that the expression levels of CD6 in double-negative (DN) thymocytes were very low and similar between WT and CD6Δd3 mice. However, there was a pronounced increase in the expression of CD6Δd3 compared with CD6WT in DP and CD4^+^ SP thymocytes, and narrower in CD8^+^ thymocytes (**Fig. 5C**). Such differences were also observed in thymocytes from adult mice, though not in splenocytes (**Fig. S4**). One possibility to explain the increase in CD6Δd3 expression is that in the absence of d3, failure of CD6 binding to its ligands could drive a compensatory mechanism to increase its expression during thymocyte development, a feature that might be elemental for maturation progression. An alternative hypothesis, though, could be conceived from the analysis of mRNA expression (**Fig. 5C**, right panel). The differences between CD6WT and CD6Δd3 protein levels do not correlate with variations in mRNA expression, as far as the levels of *CD6Δd3* transcripts are indistinguishable from those of *CD6WT* in equivalent thymocyte subsets. Thus, a possible explanation is that the presence of d3 of CD6 may be involved in the recycling/degradation of the molecule, possibly through its association with the TCR complex; in the absence of such interaction, CD6Δd3 would accumulate at the cell surface.

Of note, the levels of surface CD6 expression in WT mice also do not fully match the changes in the corresponding mRNA expression, and for example the surface CD6 levels in DP thymocytes are much lower than in CD4^+^ SP thymocytes even though the mRNA levels are identical. However, these effects in the CD6WT mice are perhaps not wholly surprising given that, especially during thymic selection, the *CD6Δd3* isoform is more prevalent in SP than in DP thymocytes (*43*). According to the above-mentioned hypothesis and considering the same rate of gene expression, an enrichment of *CD6Δd3* transcripts in CD4^+^ SP of WT mice compared with DP thymocytes would result, in the former cells, in a larger proportion of the CD6Δd3 protein being produced at the expense of CD6WT and having as consequence an increase in surface expression, due to the defect in recycling of CD6Δd3.

One direct effect of the increase in the expression levels of CD6 in thymocytes of young CD6Δd3 mice was a subtle but consistent decrease of CD5 levels in CD6Δd3 mice relatively to WT animals (**Fig. 5D**), although this compensatory effect is lost in thymocytes of adult mice (**Fig. S4A**, right panel). A compensatory increase of CD6 in CD5-KO mice has been previously described, but only on splenocytes and not on thymocytes (*36*). In contrast, in CD6-null mice it was described an actual decrease of CD5 expression in both thymocytes and splenocytes (*36*). Thus, although CD5 and CD6 may have some similar roles in T cell development and activation, it is still difficult to establish any precise reciprocal functions or how these molecules compensate for one another.

### CD6Δd3 thymocytes and mature T cells have altered responsiveness to CD3+CD28 mAb triggering

The aforementioned imbalance in thymocyte maturation suggested that the control of CD6 expression levels might be important for setting optimal levels of TCR avidity that deliver the precise signals for selection. However, these experiments did not provide direct information on the responsiveness of the developing thymocytes. Given that the transcription factor Nur77 has been correlated with the strength of TCR signaling (*50–52*), we assessed the changes in Nur77 expression in thymocytes upon TCR triggering. Thymocytes from WT and CD6Δd3 mice were activated with CD3 + CD28 mAbs and the levels of Nur77 were assessed in the different sub-populations at the indicated time points. We observed a marked decrease in Nur77 expression in CD4 and CD8 SP thymocytes from young CD6Δd3 mice, compared with the cells from WT mice, 12 h upon polyclonal activation (**Fig. 6A**), and also in CD4 SP, though not in CD8 SP thymocytes from adult mice (**Fig. 6B**). Interestingly, these decreases in Nur77 expression almost perfectly matched the converse enhanced expression levels of functionally inhibitory CD6 molecules, in exactly the same SP thymocyte populations of CD6Δd3 mice (**Fig. 5C** and **Fig. S4A**).

**Fig. 6.**
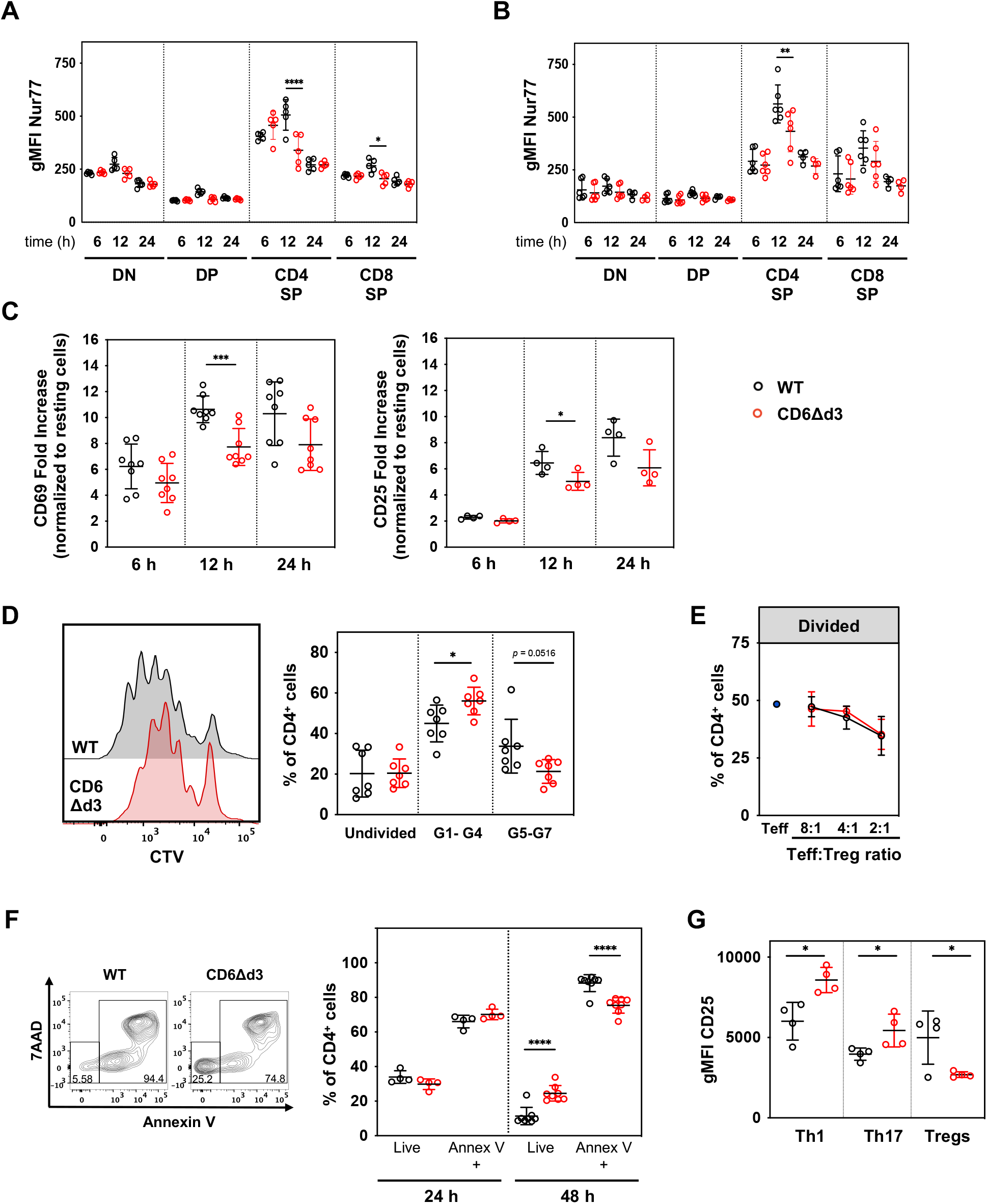
Absence of CD6d3 decreases global thymocyte and mature T cell activation induced by CD3+CD28 mAbs, but favors pro-inflammatory responses. (**A-B**) Nur77 expression upon 6, 12 and 24 h of activation with plate-bound CD3 and soluble CD28 mAbs of *ex vivo* thymocytes collected from young (A) or adult (B) WT (black circles) or CD6Δd3 (red circles) mice. (**C**) Purified CD4^+^ T splenocytes from naïve WT and CD6Δd3 mice were activated with plate-bound CD3 and soluble CD28 mAbs. T cell activation was assessed by CD69 (left) and CD25 (right) expression after 6, 12 and 24 h. Fold increase obtained by normalizing the gMFI values of activated with those of nonactivated cells. (**D**) Proliferation of purified CD4^+^ naïve splenocytes from 8 to 12-week-old WT and CD6Δd3 mice assessed upon 96 h of activation. Left, cell trace violet (CTV) dye representative histogram profiles; right, fraction of cells that did not progress from G0 (undivided), had between 1 and 4 (G1-G4), or between 5 and 7 (G5-G7) rounds of divisions. (**E**) Mouse CD4^+^ T effector (Teff) cells and Tregs were sorted based on CD25 expression. CFSE-stained Teff from WT mice were activated with CD3 mAbs and irradiated APCs in the absence (Teff) or presence of Tregs from WT or CD6Δd3 mice at different ratios. Results show the mean percentage ± SD of divided CD4^+^ T cells of four individual mice upon 72 h of culture, in one representative of two independent experiments. (**F**) Survival and apoptosis of CD4^+^ T cells after 24 and 48 h of polyclonal activation, assessed by Annexin V and 7AAD staining. Left, representative contour plot of live and annexin V positive cells after 48 h of activation; right, percentage of live and apoptotic cells after 24 and 48 h of activation. (**G**) CD4^+^ T cells from WT and CD6Δd3 mice were activated and cultured under Th1, Th17 or Treg polarization conditions for 96 h. T cell subsets were defined as Th1, CD4^+^CD25^+^IFNy^+^; Th17, CD4^+^CD25^+^IL-17^+^; Tregs, CD4^+^CD25^+^Foxp3+. The gMFI of CD25 was analyzed for each differentiated subpopulation. Each dot represents one individual mouse, horizontal lines indicate the respective mean ± SD. *, *p* < 0.05; **, *p* < 0.01; ***, *p* < 0.005; ****, *p* < 0.001, unpaired Student’s t test with or without the Welch’s correction and multiple t tests with Welch’s correction.

We next assessed the functional capacity of mature cells, isolating CD4^+^ T cells from naïve splenocytes of 8- to 12-week-old adult mice and activating them with CD3 and CD28 mAbs. We observed that CD4^+^ T cells from CD6Δd3 mice were less prone to activation than CD4^+^ T cells from WT mice, as assessed by the comparatively lower expression of CD69 and CD25 after 12 h of stimulation (**Fig. 6C**), and had fewer rounds of cell divisions (**Fig. 6D**). Mechanistically, these results were somewhat unexpected considering that the CD6WT and CD6Δd3 molecules contain the same intracellular signaling motifs/domains and that their expression levels, unlike on thymocytes, are identical in the peripheral cells of both strains (**Fig. S4B**). These differences in activation and total proliferation could not be explained by defects in the Treg compartment, given that the frequency and number of Tregs in the periphery were equivalent in WT and CD6Δd3 mice (**Fig. 4G**) and, moreover, Tregs from both mouse strains were equally functional in suppressing the activation of CD4 T^+^ effector cells from WT mice (**Fig. 6E**). Therefore, these observations indicate that on overall CD6Δd3 peripheral cells have lower reactivity than their WT counterparts in response to CD3+CD28 mAbs.

The lower responsiveness of CD6Δd3 T cells may, nonetheless, result in their increased resistance to enter apoptosis as we observed in **Fig. 6F**, resulting in a higher survival of CD6Δd3 T cells when compared with WT T cells. Also remarkably, when we activated T cells grown under Th1 or Th17 polarizing conditions, cells from CD6Δd3 mice showed increased activation of the Th1 and Th17 subsets and decreased activation of the Treg compartment (**Fig. 6G**). Taken together, these results show that WT and CD6Δd3 T cells display intrinsic differences that impact on their levels of reactivity, regardless of CD6 ligand-engagement. The combination of a higher resistance to apoptosis and an enhanced reactivity under polarizing conditions may predispose T cells from WT and CD6Δd3 to react differently in response to environmental stimuli.

### Increased susceptibility to EAE induction and amplified frequency and number of pathogenic CD4^+^ T cells in the brain of CD6Δd3 mice

Polymorphisms in the human *CD6* gene shown to increase the expression of the CD6Δd3 isoform have been associated with higher susceptibility to develop multiple sclerosis (MS) (*45, 46*). To assess whether the different CD6 isoforms display a similar correlation in mice, we induced EAE in both WT and CD6Δd3 mice by immunization with the MOG_35-55_ peptide emulsified in CFA. We found that CD6Δd3 mice developed more severe clinical manifestations together with an earlier disease onset, than WT animals (**Fig. 7A**). However, spinal cord histopathology at day 17 post EAE induction showed only negligible increases in cellular infiltration and demyelination in CD6Δd3 compared with WT mice (**Fig. 7B**). Moreover, flow cytometry analyses of brain parenchyma at different time points of disease progression did not show any differences in cellular infiltrates (CD45^high^ cells) (**Fig. 7C**), in the number or frequency of infiltrating CD4^+^ T cells or in the expression levels of the activation marker CD44 within the CD4^+^ T cell population (**Fig. 7D**), or in the number or frequencies of all other lymphoid or myeloid populations (**Fig. S5A**), when comparing WT and CD6Δd3 brains.

**Fig. 7.**
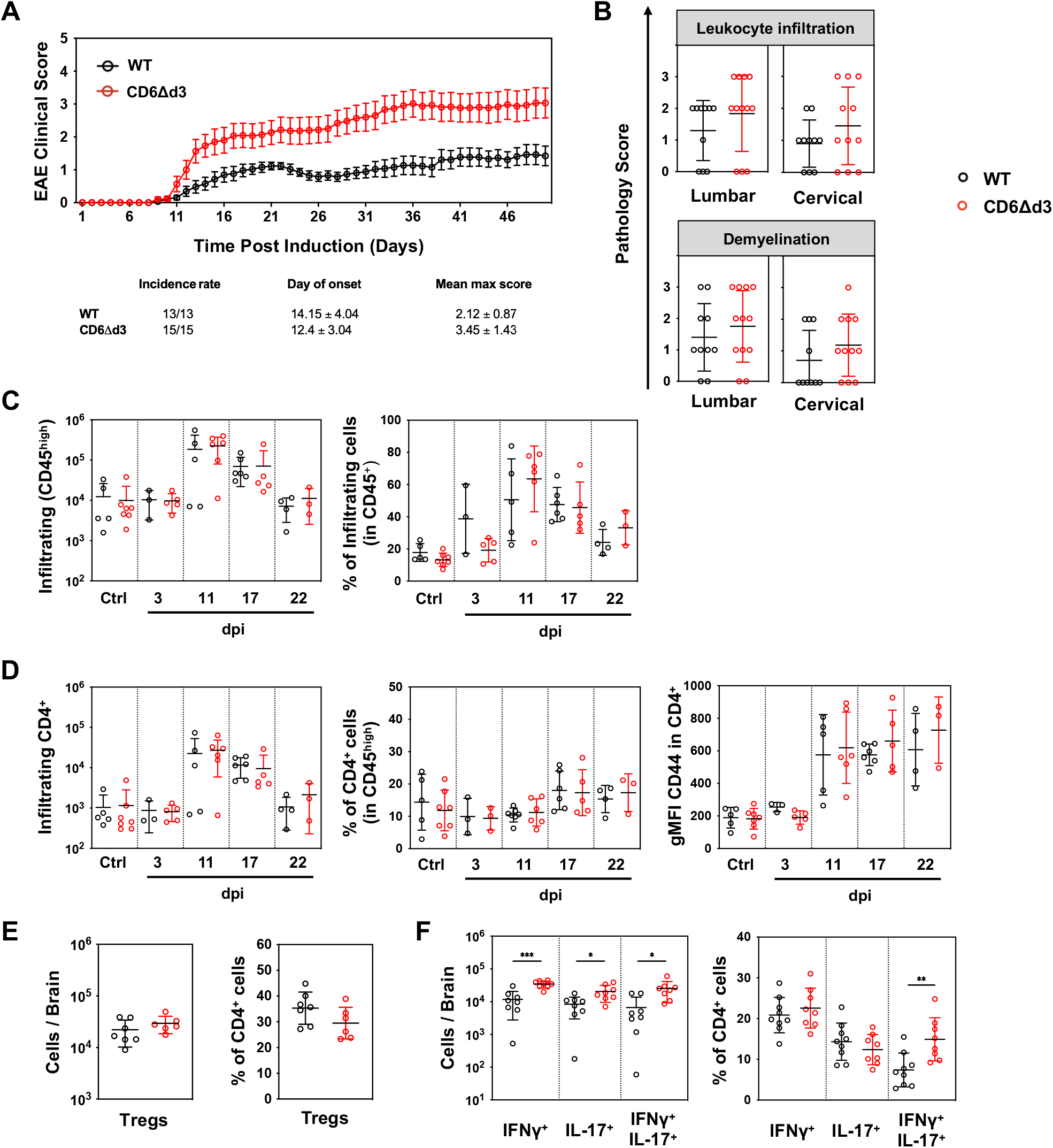
CD6Δd3 mice develop more severe EAE than WT mice. (**A**) Mean cumulative clinical EAE score from MOG_35-55_ immunized C57BL/6 WT (black circles) and CD6Δd3 (red circles) mice. Data shown are the mean ± SEM of 13 WT and 15 CD6Δd3 mice from 3 independent experiments. The differences reach statistical significance from day 28 post induction until the end of the experiment (twoway ANOVA with Sidak’s multiple comparisons test). (**B**) Pathology scores of spinal cord lumbar (left graphs) and cervical (right graphs) sections at day 17 post EAE induction. Upper panel, analysis of spinal cord leukocyte infiltration in Hematoxylin and eosin-stained sections. Lower panel, demyelination analysis by evaluation of the myelin sheet in Luxol Fast blue stained sections. Scoring: 0, no signs of leukocyte infiltration/demyelination; 1, mild leukocyte infiltration/demyelination; 2, moderate leukocyte infiltration/demyelination; 3, severe leukocyte infiltration/demyelination. (**C**) Total cell number (left) and frequency (right) of CD45^high^ cells (within all CD45^+^) infiltrating the brain parenchyma at 0 (control), 3, 11, 17 and 22 days post EAE induction, measured by flow cytometry. (**D**) CD4^+^ total cell number (left), and frequency of CD4^+^ cells (within the CD45^high^ population) infiltrating the brain parenchyma (middle) at the different days after EAE induction. Right graphs shows the gMFI for the activation marker CD44 among CD4^+^ T cells infiltrating the brain, normalized to the corresponding unstained. (**E**) Total number (left) and frequency (right) of Tregs (CD4^+^CD25^+^Foxp3^+^) in the CNS at day 17 after induction of EAE. (**F**) Cell numbers (left graph) and percentage (right graph) of IFNy-, IL-17-, and IFNy + IL-17-expressing CD4^+^ T cells isolated from the CNS at day 17 post EAE induction. (B-F) Each dot represents one individual mouse, and horizontal lines indicate the respective mean ± SD. *, *p* < 0.05; **, *p* < 0.01; ***, *p* < 0.005 (unpaired Student’s t test with Welch’s correction). dpi: days post induction.

Considering that the number of autoreactive CD4^+^ T cells reaching the brain parenchyma was comparable between WT and mutant mice, we evaluated the effector phenotype of these cells. There were no differences in the number or percentage of Tregs, one subset which could potentially contribute to lower immunosuppression favoring the development of EAE (**Fig. 7E** and **Fig. S5B**). Strikingly, we observed a significant increase in the number of IFNγ-producing (Th1), of IL-17-producing (Th17), and of IFNγ + IL-17 double-producing (IFNγ^+^IL-17^+^) subsets of CD4^+^ T cells in the brains of CD6Δd3 mice, compared with WT mice (**Fig. 7F** and **Fig. S5B**). Noteworthy, this resulted in a significant enrichment of the IFNγ^+^IL-17^+^ double-positive subset, which has been suggested to be associated with higher pathogenicity (*53*).

To assess whether the increased pathogenicity observed in the CD6Δd3 mice resulted from intrinsic properties of its T cells, we first induced EAE in both WT and CD6Δd3 mice, and then injected WT and CD6Δd3 MOG-reactive splenocytes into recipient C57BL/6 mice. The recipients that received CD6Δd3 cells displayed higher clinical scores compared with those that received CD6WT cells (**Fig. 8A**). This result is consistent with our hypothesis that CD6Δd3 T cells, including CD4^+^IFNγ^+^IL-17^+^ cells, are responsible for the enhanced pathogenicity.

**Fig. 8.**
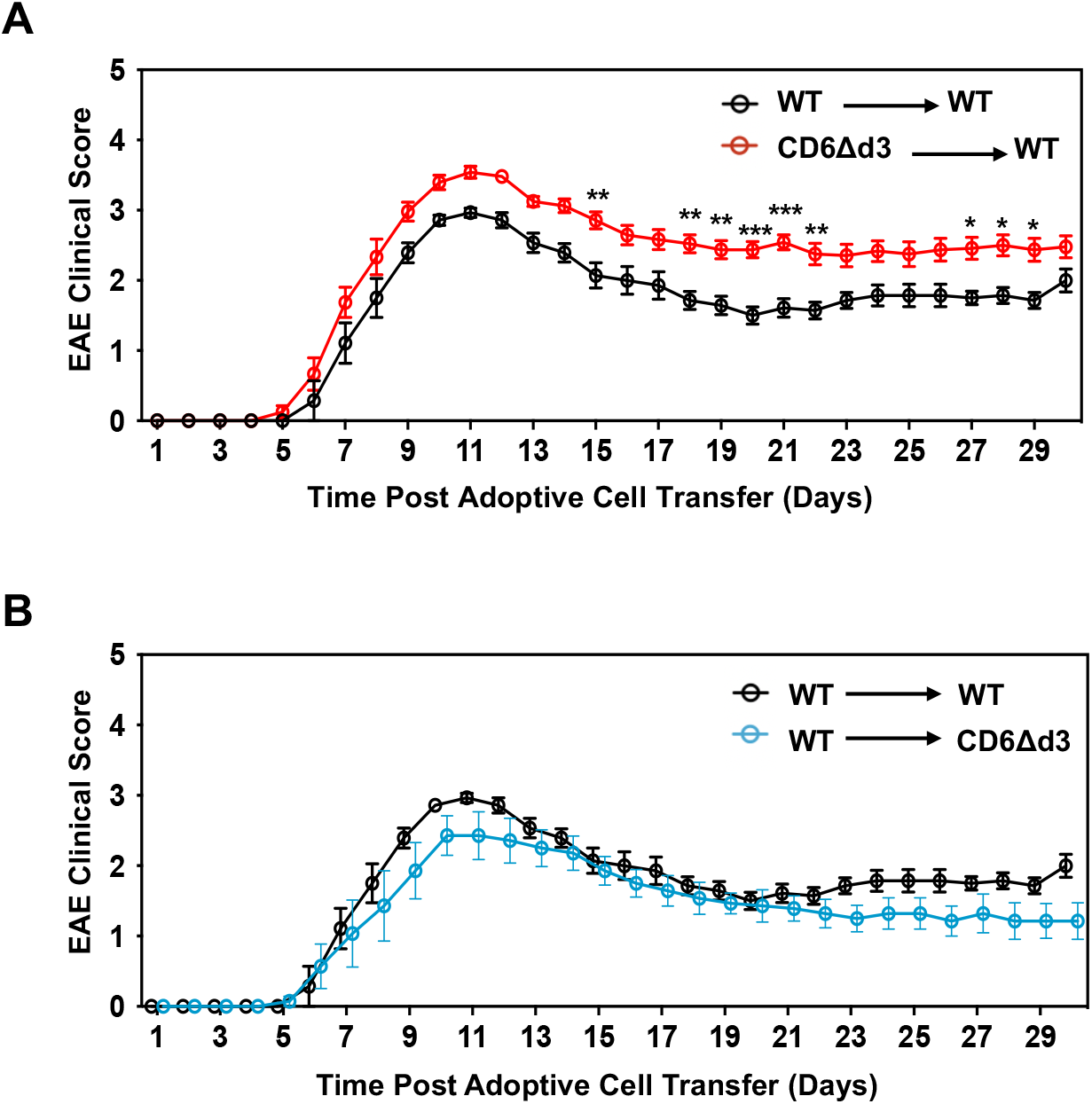
EAE susceptibility in CD6Δd3 mice is cell intrinsic. (**A**) Mean cumulative EAE clinical score of C57BL/6 mice adoptively transferred with MOG_35-55_ reactivated WT and CD6Δd3 splenocytes. Data shown are the mean ± SEM of 7 mice that received WT cells (in black circles) and 12 mice that received CD6Δd3 cells (red circles), from 2 and 3, respectively, independent experiments. (**B**) Mean (± SEM) cumulative EAE clinical score of WT (black circles) and CD6Δd3 (blue circles) mice adoptively transferred with MOG_35-55_ reactivated WT splenocytes. Data shown are representative of 2 independent experiments, with 7 mice in each group. *, *p* < 0.05; **, *p* < 0.01; ***, *p* < 0.005 (twoway ANOVA with Sidak’s multiple comparisons test).

As CD6 might interact with CD166 in the brain endothelium, we wanted to evaluate whether differences in blood-brain-barrier permeability could play a role in the increased EAE pathogenicity observed in CD6Δd3 mice. For this, we induced EAE in WT mice and then transferred CD6WT MOG-reactive splenocytes to both WT and CD6Δd3 recipients. As seen in **Fig. 8B**, no major differences were observed regarding EAE development in the WT or CD6Δd3 background, thereby excluding any major influence of the environment of CD6Δd3 mice in the increased EAE pathogenesis. Altogether, these results indicate that the different severity of EAE in WT and CD6Δd3 mice is mostly T cell intrinsic.

## DISCUSSION

It has been well understood for more than three decades that the cell surface receptor CD6 has an important role in T cell biology. For most of this time it was considered to be a costimulatory receptor. Recently, however, there has been less certainty about the nature of CD6’s contribution to T cell activation and signaling. Our previous experiments were the first to imply that in particular circumstances CD6 actually attenuates T cell activation (*25*), a finding raising the possibility that CD6 might have a dual function. In this study, we sought to reconcile the different theories of CD6 function.

A first major conclusion of the new study, consistent with our previous findings, is that the cytoplasmic domain of CD6 plays a critical role in the receptor’s inhibitory effect on T cells. Moreover, we can now also attribute to the cytoplasmic tail of CD6 a novel and critical function in its localization to the IS where it is likely to impact on the regulation of signal transduction. On the other hand, the CD6 extracellular domain may play an unanticipated role on its optimized localization, granted not by its binding to the APC-expressed ligand CD166, thus contradicting previous reports (*24, 43*), but through its interaction *in cis* with the TCR machinery.

Similarly to CD5, the nonlinear arrangement of the SRCR domains in CD6 is compatible with the establishment of interactions *in cis* with other surface molecules, including the formation of homodimers (*54–56*). Meddens *et al*. have shown that upon TCR triggering, CD6 increasingly co-localizes with the TCR/CD3 complex over time (*57*). Importantly, the artificial membranes used in their work did not include the CD6 ligand CD166. Our quantitative analysis showing the level of co-localization of CD6Δd3 with CD3 being significantly lower than that of CD6WT is thus consistent with the d3 interaction with the TCR/CD3 machinery being involved in the refined positioning of CD6 at the IS. However, it has to be taken into account that the *in vitro* models that we and others employ to establish these conclusions use Jurkat cells that express immutable forms of CD6, whereas physiological T lymphocyte-expressed CD6 cycles between the two isoforms during T cell activation. Using *ex vivo* human T lymphocytes, we have previously shown that there is an enrichment of the CD6Δd3 isoform that peaks when activation levels are at their highest, and then CD6 reverts back to the full-length form when activation is terminated (*48*). This process can be independent of APCs as shown upon PHA-mediated activation, so it is plausible that it is the specific downmodulation of d3 resulting in the uncoupling of CD6 from the TCR machinery which facilitates the progression of T cell activation.

A general principle emerging from our work and other published data is that the function of CD6 and, in particular, the intensity of the inhibitory signals it generates, is highly context dependent. Analyses of CD6-null mutant mice might, therefore, give ambiguous answers since the complete absence of the molecule could impact the biological functions of CD6 in different ways, depending on the levels of coincident signaling. Our analysis of the CD6Δd3 mice in which only d3-dependent binding functions are abrogated has helped to clarify some of the ligand(s)-binding effects of CD6.

It was suggested that the involvement of CD6 in the development or progression of neuroinflammation results from its binding to CD166 and/or to an alternative ligand, CD318, within the blood-brain-barrier, which would constitute a checkpoint for the entry of T cells into the CNS. CD166 is highly expressed in the endothelium (*35*) and CD318, although not constitutively expressed, is upregulated in brain microvascular endothelial cells upon IFNγ stimulation (*58*). Both CD6^-/-^ and CD318^-/-^ mice have been reported to have decreased T cell infiltration in the CNS, implying that a CD6-ligand interaction would be required for T cell extravasation and initiation of EAE (*35, 58*). However, in complete contrast with the full protection of CD6^-/-^ mice reported by Li *et al*. (*35*), we found that CD6Δd3 mice developed more severe EAE than the C57BL/6 CD6^+/+^ littermates. The aggravated condition occurred despite there being no major differences in leukocyte infiltration in the spinal cord or in the number of invasive leukocytes in the brain parenchyma between WT and CD6Δd3 mice. This argues against T lymphocyte infiltration into the CNS being controlled by the interaction between T cell CD6 with CD166 expressed in endothelial cells. And although it cannot be precluded a possible CNS entry checkpoint being imposed by a putative CD6-CD318 ligation, this particular interaction can occur indifferently in both WT and CD6Δd3 mice given that the interaction is established by the SRCR domain 1 of CD6. Therefore, a possible simple explanation for the exacerbated neuroinflammation in CD6Δd3 mice, compared with the WT controls, is thus the observed disproportionate frequency in the brain parenchyma of T cell subsets associated with higher pathogenicity, namely encephalitogenic IFNγ^+^/IL-17^+^ CD4^+^ T cells (*53*).

Our study of thymocyte development suggests an impaired transition to SP stages in CD6Δd3 mice, likely altering the TCR repertoire, as previously described for CD6^-/-^ mice (*36*). Given the adhesion-promoting properties of the CD6/CD166 pair, it is likely that by abolishing binding, the functional avidities of MHC-antigen complexes of thymic APCs and thymocytes are compromised, as previously suggested (*49*), resulting in defective or biased selection. Similar phenotypes have been described for other adaptors, namely LAT (*59*) and LCP2 (*60*). Interestingly, we could not observe major differences in peripheral leukocyte subset profile in CD6Δd3 mice relative to WT controls, suggesting that putative repertoire changes remain hidden until cells are exposed to a specific stimulus or environment provided, *e.g*., by the CNS, where the pathogenic potential of particular T cell subsets might be unleashed, as we observed in the CD6Δd3 mutant mice.

Besides the imbalances in the frequency of specific sub-types of T cells, such as IFNγ- and/or IL-17-producing CD4 T cells, the increase in EAE severity observed in CD6Δd3 mice could be partly due to an overall malfunctioning T cell population. An intriguing question relates to the difference in reactivity between similar populations from the WT and CD6Δd3 mutant mice, observed after polyclonal activation with CD3 + CD28 mAbs. As argued above, CD6WT and CD6Δd3 molecules would, in principle, induce equivalent levels of activation given that the intracellular signaling machinery remains identical. However, we observed that purified mouse CD6Δd3 CD4^+^ T cells were less activated and had fewer rounds of divisions than WT CD4^+^ T cells. Although perhaps counterintuitive, this effect very much recapitulates what was observed with human CD4^+^ T cells carrying the CD6 MS susceptibility allele, that display impaired proliferation upon CD3 + CD28 activation when compared with CD4^+^ T cells that carry the non-risk allele (*46*). It remains to be established whether the differences in reactivity between T cells in animal models or from MS patients provoked by *in vitro* stimulation using CD3 + CD28 mAbs reflects their physiological behavior and responses during EAE or MS.

While we have detailed, in our Jurkat cell model, the function of the different parts of the molecule, it is not possible to fully establish a parallel correlative function of these domains in the mouse models simply because CD6 in WT mice cycles between full-length and Δd3 isoforms, and on activation a large proportion of CD6 molecules may factually be CD6Δd3. Therefore, whatever signaling or adhesion functions are triggered or inhibited by the ligation of domain 3 of CD6, these are possibly relevant mostly in the initial stages of activation, and also in the termination phase when transient CD6Δd3 molecules revert back to the conventional three-SRCR domain-containing form.

We propose that CD6 represents a new class of ‘two-speed’ inhibitory receptor that regulates immune responsiveness through an internal switch mechanism. First, it sets signaling thresholds via tonic inhibitory signaling, functioning as a scaffold. But distinctively from other inhibitory receptors, CD6 can alleviate repression not because of changes in the availability of extracellular *cis* or *trans* ligands but by an alternative splicing-mediated shift to an isoform that is segregated away from the TCR machinery and thus unable to inhibit signaling, acting in this case as an immune checkpoint. Through this resourceful and intricate array of mechanisms and interactions, regulatable during the course of activation, CD6 is able to optimally tune signaling in a variety of different ways in many diverse T cell subsets.

## MATERIALS AND METHODS

### Ethics statements

All procedures involving mice were done in strict accordance with recommendations of the European Union Directive 2010/63/EU and previously approved by the competent national authority Direcção Geral de Alimentação e Veterinária (DGAV, approval no. 009951), and by the i3S Animal Welfare and Ethics Body. Animals were housed in a facility accredited by AAALAC International under specific pathogen-free conditions.

### Cell culture, CD6 constructs and lentiviral transduction

Jurkat E6.1, both parental and CD6-transfected, and Raji cell lines were grown in RPMI-1640 culture media supplemented with 10 % fetal bovine serum (FBS), 1 mM sodium pyruvate, 2 mM L-glutamine, penicillin G (50 U/ml) and streptomycin (50 μg/ml). All cell lines were maintained at 37 °C and 5 % CO2.

WT CD6 and isoform-encoding sequences were amplified by PCR from pEGFP-N1/CD6FL (*43*) by removing different segments of CD6 or point mutating single residues according to the annotated sequence NM_006725 (Genbank, NCBI), using specific primers (**Table S1**). The cytoplasmic deletion mutants, CD6-D571_stop_ and CD6-D485_stop_, have the numbered aa substituted by a stop codon, and are 570 and 484 aa-long, respectively. The CD6-tailess mutant, CD6Δcyt, still contains 6 aa of the cytoplasmic tail (K429 substituted by a stop codon), for proper membrane attachment and stability. The dsDNA composed of CD6 coding sequence where cytoplasmic tyrosines were substituted by phenylalanines was synthesized (IDT). All CD6 sequences were cloned in the lentiviral expression vector pHR using *MluI* and *NotI* restriction sites and transduced in Jurkat E6.1 and Je6-NF-κB::eGFP cell lines, as described (*61*). All the transfected cells were checked by flow cytometry for CD6 expression level homogeneity and, when needed, cell sorting was performed.

### Screening of NF-κB signaling in Jurkat reporter cell lines

The Jurkat Je6 NF-κB::eGFP reporter cell line was described previously (*62*). 5 x 10^4^ reporter cells per well were co-cultivated for 24 h with 5 x 10^4^ Raji-CD166^+^ or Raji-CD166^neg^ cells, previously loaded, or not, for 1 h with 1 μg/ml of the sAg staphylococcal enterotoxin E (SEE) (Toxin Technologies). After 24 h of culture at 37 °C, cells were harvested and the Raji cells were stained with an APC- or PE-cy7-conjugated mouse anti-human CD19 mAb, used to gate out the Raji cells during analysis. Subsequently, expression of reporter eGFP was measured by flow cytometry. Geometric mean of fluorescence intensity (gMFI) of reporter cells, excluding the CD19^+^ Raji cells, was used for further analysis. The activation index was calculated as eGFP values of cells interacting in the presence of sAg over the eGFP values of the interactions without sAg.

### Cell-cell adhesion

Jurkat E6.1 cells expressing CD6 variants were stained with 0.25 μM of the cell tracker green CMFDA (Thermo Fisher Scientific) for 20 min at 37 °C. Cells were washed and put in contact for 20 min with Raji-CD166^+^ or Raji-CD166^neg^, previously loaded, or not, for 30 min with 5 μg/ml of SEE. The interaction was stopped by the addition of 4% paraformaldehyde (PFA) for 10 min. The proportion of E6.1 and Raji is 1:1 and they interacted in a final volume of 20 μl of RPMI without FBS. Cell conjugates were analyzed by imaging flow cytometry (Amnis ImageStream) by acquiring 20,000 cells per condition and then calculating the percentage of doublets (with at least one E6.1 Jurkat-green stained cell in contact with a Raji-colorless cell) within all the stained cells.

### Immunological synapse formation, immunostaining and colocalization analysis

Raji-CD166^+^ or Raji-CD166^neg^ were incubated with 1 μg/ml of SEE and plated on poly-L-lysine-coated glass coverslips for 30 min at 37 °C. Jurkat E6.1 cells expressing the different CD6 mutants were added to the Raji cells and allowed to interact for 5 or 30 min at 37 °C. Cellular conjugates were fixed for 10 min with PFA 4%, washed and blocked with PBS-0.5% bovine serum albumin (BSA). After blocking, cells were stained sequentially with mouse anti-human CD6 (MEM98, EXBIO), secondary antibody donkey anti-mouse Alexa Fluor 568 (Thermo Fisher) and mouse anti-human CD3 (UCHT1) Alexa Fluor 488-conjugated (BioLegend). All antibody dilutions were done in blocking solution. DAPI was used for nuclear staining. Images were acquired in a Leica SP5 confocal microscope. Conjugate formation and synapse localization of CD6 were quantified with blind scoring for a minimum of 40 conjugates in each condition by three examiners.

For CD6/CD3 colocalization analysis, a series of Z-stack images were captured and the Pearson’s correlation coefficient for colocalization at the synaptic region was performed by the JACoP plugin for ImageJ (*63*). The colocalization was evaluated in 15 to 16 conjugates per condition from three independent experiments.

### Development of CD6Δd3 mice and immunophenotyping

The CD6Δd3 mutant C57BL/6 mouse was generated by pronuclear injection of a mix containing two sgRNAs (2.5 ng/μl each), targeting the sequences 5’-GGGCAGGATATGTTTACCAG-3’ and 5’-ATTTTGTCTCTGCCGACAGC-3’, located respectively in the upstream and downstream introns flanking exon 5, Cas9 mRNA (5 ng/μl), and the replacement ssDNA oligonucleotide 5’-GAGACCAGTACTGTGGTCACAAGGAGGACGCAGGAGTGGTGTGCTCAGGTCAGTGAGCCTGGAGGGCAGGATATGTTTACAGCTGGCATATACTGGCCCAAGAAAGACCATTGCACTTACCATGCACAGGCAGACATTAATTAGGATGGTTTTGTTTTTT-3’ (10 ng/μl) (obtained from IDT), in 10 mM Tris-HCl pH7.5; 0.1 mM EDTA. The sgRNAs were generated by annealing oligos 5’-AGGGGGGCAGGATATGTTTACCAG-3’ with 5’-AAACCTGGTAAACATATCCTGCCC3’ and 5’-AGGGATTTTGTCTCTGCCGACAGC-3’ with 5’-AAACGCTGTCGGCAGAGACAAAAT-3’, and introducing them into the plasmid gRNA basic (*64*). The plasmids were linearized and the sgRNAs synthesized by in vitro transcription (Megashort Script in vitro T7, Ambion), purified (Megaclear RNA kit, Ambion) and analyzed on Experion (Bio-Rad) for integrity, purity and yield check.

In-depth immune-phenotyping analysis was performed on spleens and thymi of six 12-week-old C57BL/6/N and CD6Δd3 mice (3 males, 3 females per group) by multi-parametric flow cytometry. Leukocytes were isolated from thymi and spleens by combined mechanical treatment and enzymatic digestion using a GentleMACS™ octo dissociator system (Miltenyi). Cell suspensions were numerated for all samples on AttuneNxT flow cytometer. Erythrocyte were lysed using RBC Lysis solution (eBioscience, #00-4333-57) following manufacturer recommendations. Fc-receptors were blocked using anti-CD16/CD32 (#24G2) hybridoma. Cells were stained in one step with corresponding antibody panel (**Table S2**), washed and analyzed within 4 h after organ collection. Dead cells were removed using Live/Dead SytoxBlue staining (Thermo, #S11348). Datasets were acquired on a 5-laser BD LSRFortessa™ SORP cell analyzer using an HTS plate reader on a standardized analysis matrix using BD FACSDiva 8.0.1 software. Data were visualized using radar plot on Excel or Tableau Desktop software. Frequencies of cell subsets on all mice of the study were pooled in one single spreadsheet per panel per organ, transformed in asinh and centered to the mean. The resulting variable list was run in SIMCA Multivariate analysis software (Sartorius). First, distribution of samples in PCA score was considered to remove potential outliers. Next, an OPLS-DA method was applied on the remaining dataset to identify groups of samples presenting similar types of variations. Overall, the distribution of mice was similar before and after OPLS-DA modelling for each experimental group. Variables important for this projection (VIP) were selected with a VIPpred value over 1. A final PCA was run on data only coming from these selected predictive variables to ensure that the OPLS-DA model was not overfitting the dataset. List of phenotypes with VIP>1 were further visualized on moustache plots using Tableau Desktop software.

For conventional immunophenotyping, bone marrow cells, splenocytes and thymocytes were collected and stained with diverse antibody panels. Mix 1 for spleen and bone marrow, CD45 PerCP/Cy5.5 (I3/2.3), CD19 APC (6D5), CD3 APC/Cy7 (17A2), CD4 FITC (RM4-5), CD8 Pacific Blue (53-6.7), CD6 PE (BX222) and CD5 PE/Cy7 (53-7.3); mix 2 for spleen, CD45 PerCP/Cy5.5, CD3 APC/Cy7, CD4 FITC, CD8 PE, CD44 PE/Cy7 (IM7) and CD62L APC (MEL-14); mix 3 for spleen, CD45 PerCP/Cy5.5, CD3 APC/Cy7, CD4 FITC, CD25 PE (PC61) and Foxp3 APC (150D) after cell fixation and permeabilization (fixation/permeabilization kit, eBiosciences). Mix 1 for thymus, CD45 PerCP/Cy5.5, CD3 APC/Cy7, CD4 FITC, CD8 Pacific Blue, CD6 PE, CD5 PE/Cy7; mix 2 for thymus, CD45 PerCP/Cy5.5, CD4 FITC, CD25 PE and intranuclear staining with Foxp3 APC. For analysis of young thymi, thymocytes from mice with 1 week were stained with CD3 APC/Cy7, CD4 FITC, CD8 PE, CD44 APC, CD25 PerCP/Cy5.5, CD69 BV421 (H1.2F3) and intranuclear staining with Foxp3 APC. All the above-mentioned antibodies are from BioLegend. Data were acquired in a FACSCanto II flow cytometer (BD Biosciences). Post-acquisition analysis was performed using FlowJo software v10 (Tree Star).

### qRT-PCR

Double negative, double positive, CD4^+^ and CD8^+^ thymocytes were obtained by cell sorting (FACS Aria, BD Biosciences) from one-week-old mice. RNA was extracted using TRIzol (Invitrogen). cDNA was synthesized using Superscript IV reverse transcriptase (Invitrogen) and random hexamers (Merck). qRT-PCR reactions were performed using iTaq Universal SYBR Green Supermix (Bio-Rad) and using the following oligos for mouse *Cd5* (forward, 5’-CCATGGACTCCCACGAAGTG-3’; reverse, 5’-GCCACTTAGCATCACCTGGA-3’); mouse *Cd6* (forward, 5’-GTGCAGGACCAGGAGTGTAG-3’; reverse, 5’-ACCATCCACTAACCGCACTG-3’) and mouse hypoxanthine-guanine phosphoribosyltransferase gene (*Hprt*) (forward, 5’-GGACTTGAATCAAGTTTGTG-3’; reverse, 5’-CAGATGTTTCCAAACTCAAC-3’) in a CFX96 Real-Time PCR Detection System (Bio-Rad). The qRT-PCR results were analyzed using the ΔCt method (relative expression) (*65*), using *Hprt* as the reference gene.

### *In vitro* T cell activation, proliferation and apoptosis assays in CD4^+^ T cells

CD4^+^ T cells were isolated from naïve gender- and age-matched WT and CD6Δd3 splenocytes by negative selection using magnetic beads (Mojo sort kit, BioLegend). Isolated cells were activated with 3 μg plate-bound anti-mouse CD3 (145-2C11) and 5 μg soluble anti-CD28 (37.51), both from BioLegend. For activation assays, upregulation of CD69 and CD25 was assessed by flow cytometry 6, 12 and 24 h later. Fold increase was obtained by normalizing the gMFI of CD69 Pacific Blue or CD25 PE upon activation with the respective unstimulated values. For apoptosis assays, CD4^+^ T cells were allowed to activate for 24 and 48 h and then stained with Annexin V FITC and 7AAD (BD Pharmingen). Cell proliferation was analyzed 96 hours post polyclonal activation in CD4^+^ T cells previously stained with cell trace violet dye (Thermo Fisher).

### Nur77 transcription factor expression in activated thymocytes

Thymocytes from one-week-old and adult (8-12 weeks) mice were collected and cultured in 96-well plates, previously coated with 3 μg anti-mouse CD3 (145-2C11) monoclonal antibody, in the presence of 5 μg soluble anti-CD28 (37.51). Nur77 expression was measured by flow cytometry upon 6, 12 and 24 h of activation. Cells were stained with CD4 BV510, CD8 Pacific Blue, CD25 FITC, CD69 PE/Cy7, and intranuclearly with Nur77 PE (12.14), all from Biolegend.

### *In vitro* Th1, Th17 and Treg polarization

Following the isolation and activation procedures described above, for Th1 polarization the medium was supplemented with 5 ng/ml IL-2 and 10 ng/ml IL-12 (both from Peprotech), and 10 μg/ml anti-IL-4 (11B11, a kind gift from Luís Graga, IMM, Lisbon). For Th17 polarization, the medium included 1 ng/ml TGF-β (R&D), 10 ng/ml IL-1β and 20 ng/ml IL-6 (both from Peprotech), and 10 μg/ml anti-IFNγ (R46A2, gently given by Luís Graga). Treg polarization was induced with the supplementation of 5 ng/ml IL-2 and 5 ng/ml TGF-β.

### EAE induction and disease score assessment

Active EAE was induced in 8- to 12-week-old female mice by inguinal *s.c*. immunization with 200 μg MOG_35-55_ (MEVGWYRSPFSRVVHLYRNGK) peptide (Schafer-N) emulsified in complete Freund’s adjuvant (CFA) suspension (4 mg/ml *Mycobacterium tuberculosis*) (Mdbiosciences) and *i.v*. injection of 200 ng pertussis toxin (PTX) (Quadratech Diagnostics) on days 0 and 2 following immunization. Clinical severity was assessed daily with a 0-5 scoring system: 0, no signs; 1, tail atony; 2, hind limb weakness; 3, partial hind limb paralysis; 3.5, flattening of hind quarters with complete paralysis; 4, partial front paralysis; 4.5, quadriplegia; 5, moribund or dead.

### Adoptive transfer EAE induction

Active EAE was induced in donor mice as described above but without PTX injection. On day 11, these mice were killed and splenocytes were collected. Isolated cells were cultured for 72 h in complete RPMI further supplemented with nonessential amino acids, β-mercaptoethanol, 20 ng/ml of IL-12 (Peprotech), 10 μg/ml anti-IFNγ (XMG1.2, BioLegend) and 20 μg/ml MOG_35-55_. On the day of recipient mouse injection, cells were harvested, washed and counted. 25 x 10^6^ cells were injected *i.p*. in each recipient female mouse together with *i.v*. injection of 200 ng PTX on days 0 and 2 following cell transfer. The scoring system used was as described above.

### Central nervous system histopathology and flow cytometry

Mice were deeply anesthetized at indicated time points for transcardiac perfusion with cold PBS. For histopathology examination, spinal cords were immersed in neutral buffered formalin for 24 h and were then removed from the bone and fixed for an extra 24 h before being routinely processed and embedded in paraffin. Histologic 4 μm-thick sections were obtained for staining. Hematoxylin and eosin (H&E) and Luxol fast blue (LFB) staining were performed according to per standard protocols. Pathology scores for leukocyte infiltration and demyelination were performed by an experienced pathologist, in a blind fashion, and then validated by two other independent examiners. A 0-3 scoring system was used: 0, no signs of leukocyte infiltration/demyelination; 1, mild leukocyte infiltration/demyelination; 2, moderate leukocyte infiltration/demyelination; 3, severe leukocyte infiltration/demyelination.

For flow cytometry of brain tissue, brains were excised, gently dissociated, and cells were passed through a 100 μm cell strainer. Stock isotonic Percoll (GE Healthcare) (SIP) 90% (in HBSS without Ca^2+^ and Mg^2+^) was added to each cell suspension to obtain a final 30% SIP. Slowly, the cell suspension was added on top of the 70% SIP avoiding mixing the 70 and 30% solutions. Percoll gradients were centrifuged (500 x *g*, 30 min, at 22 °C) and the enriched population of leukocytes/microglia was collected at the 70–30% interphase. After isolation, cells were washed and counted. Thereafter, 5 x 10^5^ cells were incubated with CD3 APC/Cy7 (17A2), CD4 FITC (RM4-5), CD19 APC (6D5), Ly6G Pacific Blue (1A8), CD45 PerCP/Cy5.5 (I3/2.3), CD11b PE (M1/70), CD44 PE/Cy7 (IM7), all from Biolegend.

For intracellular staining of mouse cytokines, after the density gradient protocol cells were activated for 5 h with 500 ng/ml ionomycin and 50 ng/ml phorbol 12-myristate 13-acetate (PMA), in the presence of 10 μg/ml brefeldin A (BFA), all from Sigma-Aldrich. After staining for surface antigens with CD45 PerCP/Cy5.5 (I3/2.3), CD11b PE (M1/70), CD4 FITC (RM4-5) and TCRβ eFluor 450 (H57-597, from eBioscience), cells were fixed and permeabilized with the eBioscience fixation/permeabilization kit. Intracellular staining with IL-17A APC (TC11-18H10.1) and IFNγ PE (XMG1.2), from BioLegend.

For Treg specific staining, after the density gradient protocol, cells were stained with CD45 PerCP/Cy5.5 (I3/2.3), CD11b PE (M1/70), CD4 FITC (RM4-5), TCRβ eFluor 450 (H57-597) and CD25 PE (PC61, BioLegend), fixed and permeabilized as described above, and intranuclear stained for Foxp3 (150D, BioLegend).

Data were acquired in a FACSCanto II flow cytometer (BD Biosciences). Post-acquisition analysis was performed using FlowJo software v10 (Tree Star).

### Statistical analysis

All data were analyzed with the GraphPad Prism software (v.7, GraphPad software Inc. CA). Results are presented as means ± standard deviation (SD) or ± standard errors of the means (SEM). Specific statistic tests are discriminated in each figure legend. Only *p*-values < 0.05 were considered statistically significant (*, *p* < 0.05; **, *p* < 0.01; ***, *p* < 0.005; ****, *p* < 0.001).

## Acknowledgements

We thank M Brown (University of Oxford) for helpful discussions and review of the manuscript. The authors acknowledge the support of the i3S scientific platforms Advanced Light Microscopy (ALM), Bioimaging, member of the national infrastructure PPBI - Portuguese Platform of Bioimaging (PPBI-POCI-01-0145-FEDER-022122), Translational Cytometry, and Animal Facility.

