## Supplementary Figures for "The dual character of the inhibitory functions of CD6"

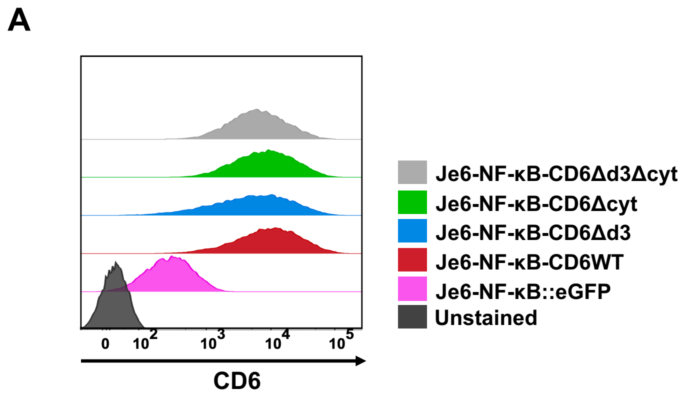


**Fig. S1. Expression of CD6 WT and mutants in Je6-reporter cells.** Flow cytometry analysis of MEM98 (anti-CD6 mAb) labeling of Je6-NF-κB::eGFP reporter cells expressing CD6WT, CD6Δd3, CD6Δcyt or CD6Δd3Δcyt.

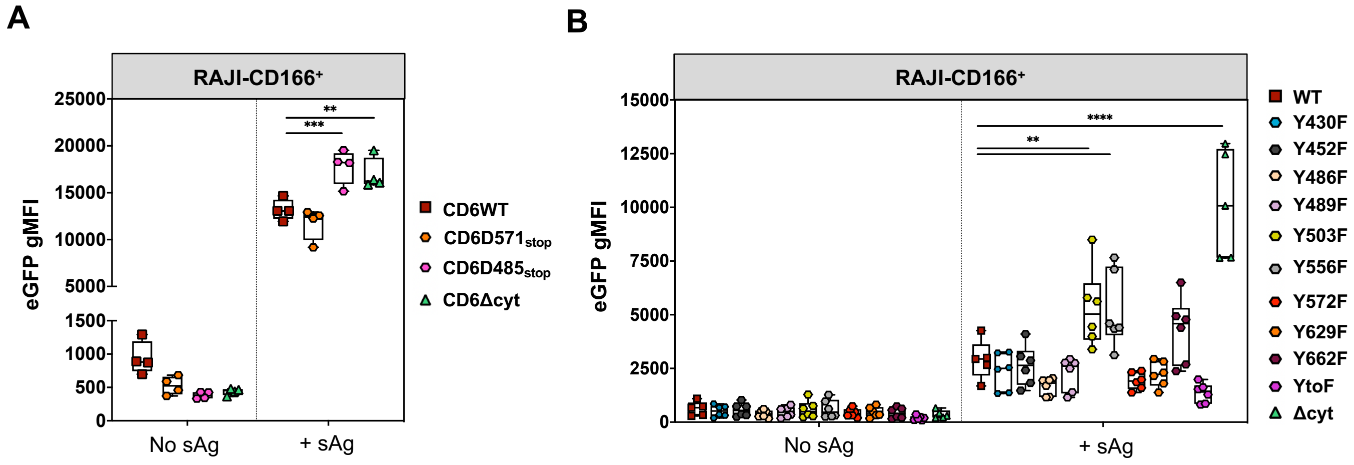


**Fig. S2. CD6 cytoplasmic tail sequences and motifs impact on T cell activation.** (**A**) Je6-NF-κB::eGFP cells expressing different CD6 cytoplasmic tail truncations were allowed to interact for 24 h at 37 ºC with Raji-CD166+ cells, presenting or not sAg. NF-κB-eGFP upregulation was assessed by flow cytometry. Each dot represents one individual experiment performed with technical duplicates. Results showing the gMFI for the conditions without and with sAg. (**B**) Je6-NF-κB::eGFP cells expressing different CD6 cytoplasmic tyrosine-to-phenylalanine substitutions were co-cultured for 24 h at 37 ºC with Raji-CD166+ cells, presenting or not sAg. NF-κB-eGFP up-regulation was assessed by flow cytometry. Graphs show the gMFI for the conditions without and with sAg for tyrosine single substitutions and full Y-to-F substitutions. Statistical analysis relative to the respective CD6WT, **, $p < 0.01$; ***, $p < 0.005$; ****, $p < 0.001$, two-way ANOVA, followed by Turkey's multiple comparison test.

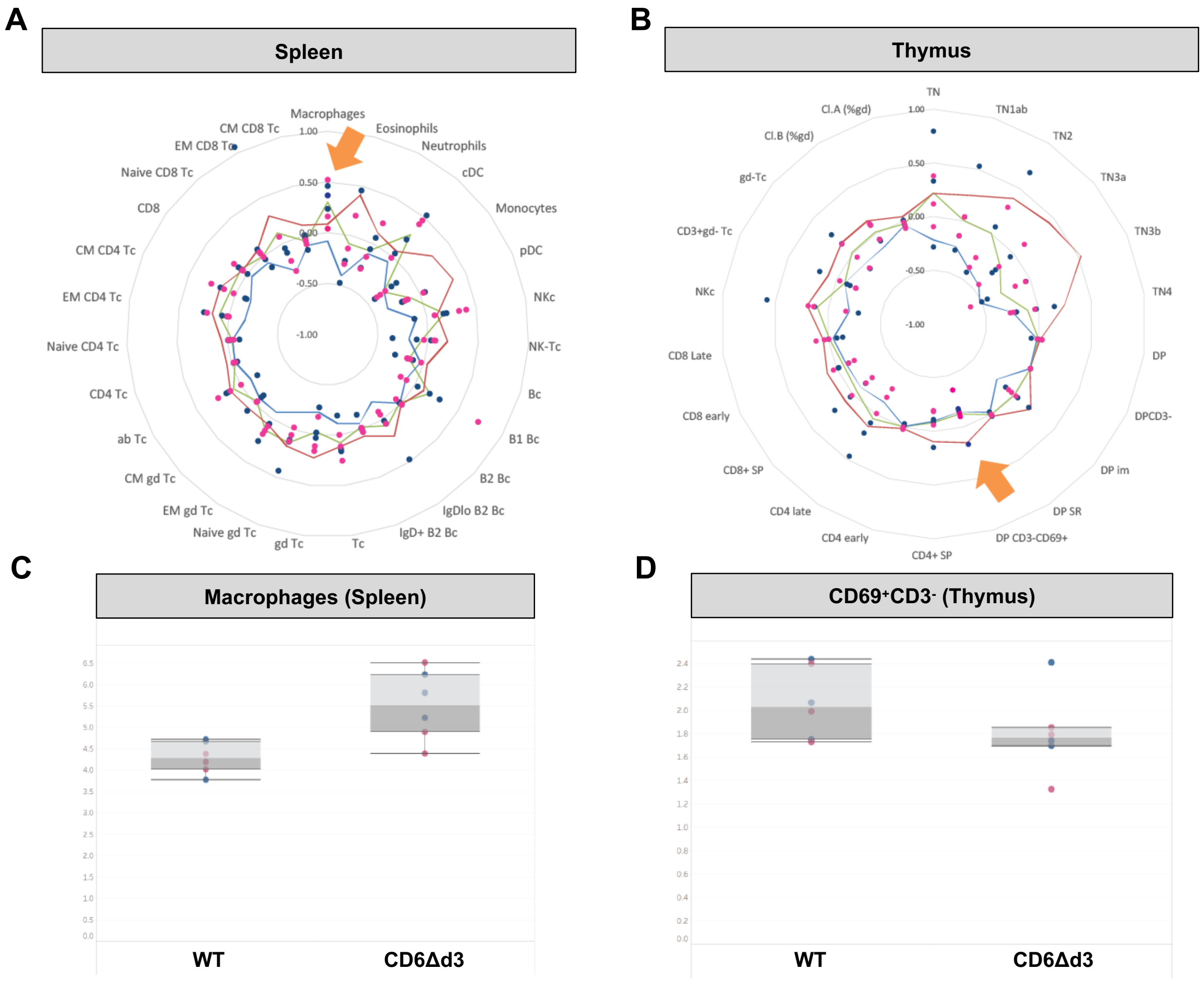


**Fig. S3. FC Radar plot of leukocyte proportion variations between CD6Δd3 and control animals in spleen (A) and thymus (B).** (**A-B**) EM: effector memory ($CD44^{+}CD62L^{-}$); CM: central memory ($CD44^{+}CD62L^{+}$); naïve ($CD44^{-}CD62L^{+}$); TN: triple negative ($CD3^{-}CD4^{-}CD8^{-}$); DP: double positive ($CD4^{+}CD8^{+}$); DP im: immature DP ($CD4^{+}CD8^{+}CD69^{-}CD71^{+}CD3^{-}$); DP SR: DP small resting ($CD4^{+}CD8+CD69^{-}CD71^{-}CD3^{-}$). Filled circles: individual values of treated animals; green: average variation of subset proportion in CD6Δd3 animals compared with WT animals [(Mean CD6Δd3)/(Mean WT)-1]; blue: lower limit of WT value dispersion [(Mean WT - 1 SD)/(Mean WT)-1]. Each circle (blue: male; pink: female) represents a value obtained in independent WT mice (value measured in (CD6Δd3n)/(Mean WT)-1). Values expressed in asinh ratio. This high content analysis exemplified the absence of major leukocyte changes in CD6Δd3 mice besides those depicted by orange arrows, in which variation in CD6Δd3 mice is either increased (macrophages) or decreased ($CD69^{+}CD3^{-}$ DP) compared with WT mice. (**C**) Proportion of macrophages in spleen differs between WT and CD6Δd3 mice. (**D**) Proportion of $CD69^{+}$ thymocytes among $CD3^{-}$ DP cells is decreased in CD6Δd3 mice compared with WT mice.

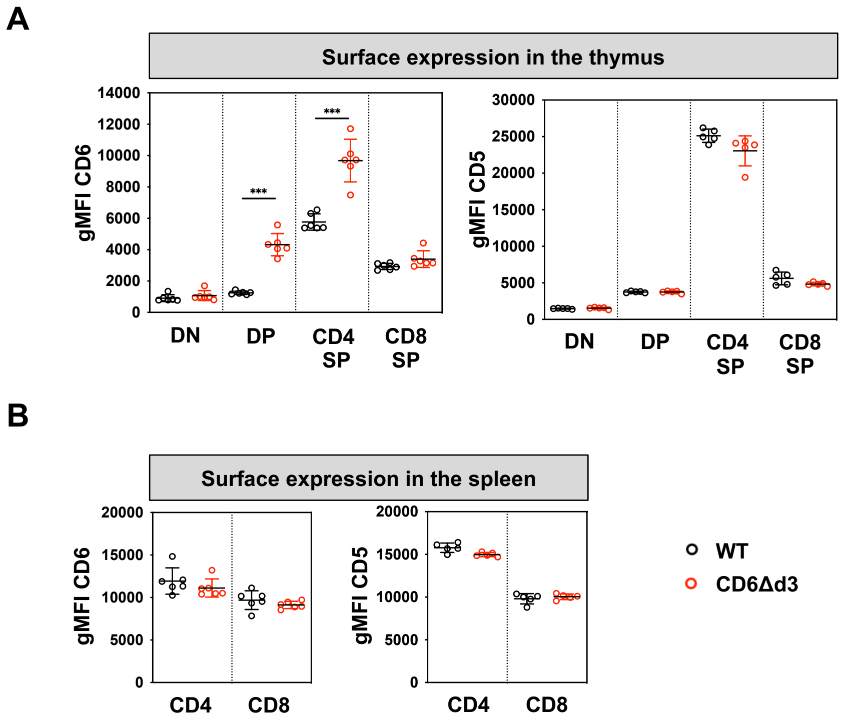


**Fig. S4. CD6 and CD5 expression in adult WT and CD6Δd3 mice.** (**A**) Surface expression of CD6 (left) (staining with anti-CD6d1 mAb BX222) and CD5 (right) in the different thymocyte subsets (DN, DP, $CD4^+$ SP and $CD8^+$ SP T cells) of WT (black) and CD6Δd3 (red) adult mice (8 – 12-weeks old), measured by flow cytometry. (**B**) Surface expression of CD6 (left) and CD5 (right) in splenocytes from adult mice. DN: double negative; DP: double positive; SP: single positive. Each dot represents one individual mouse, and the results show the mean ± SD. ***, $p < 0.005$, unpaired Student's t test with Welch's correction. Each experiment was performed at least twice.

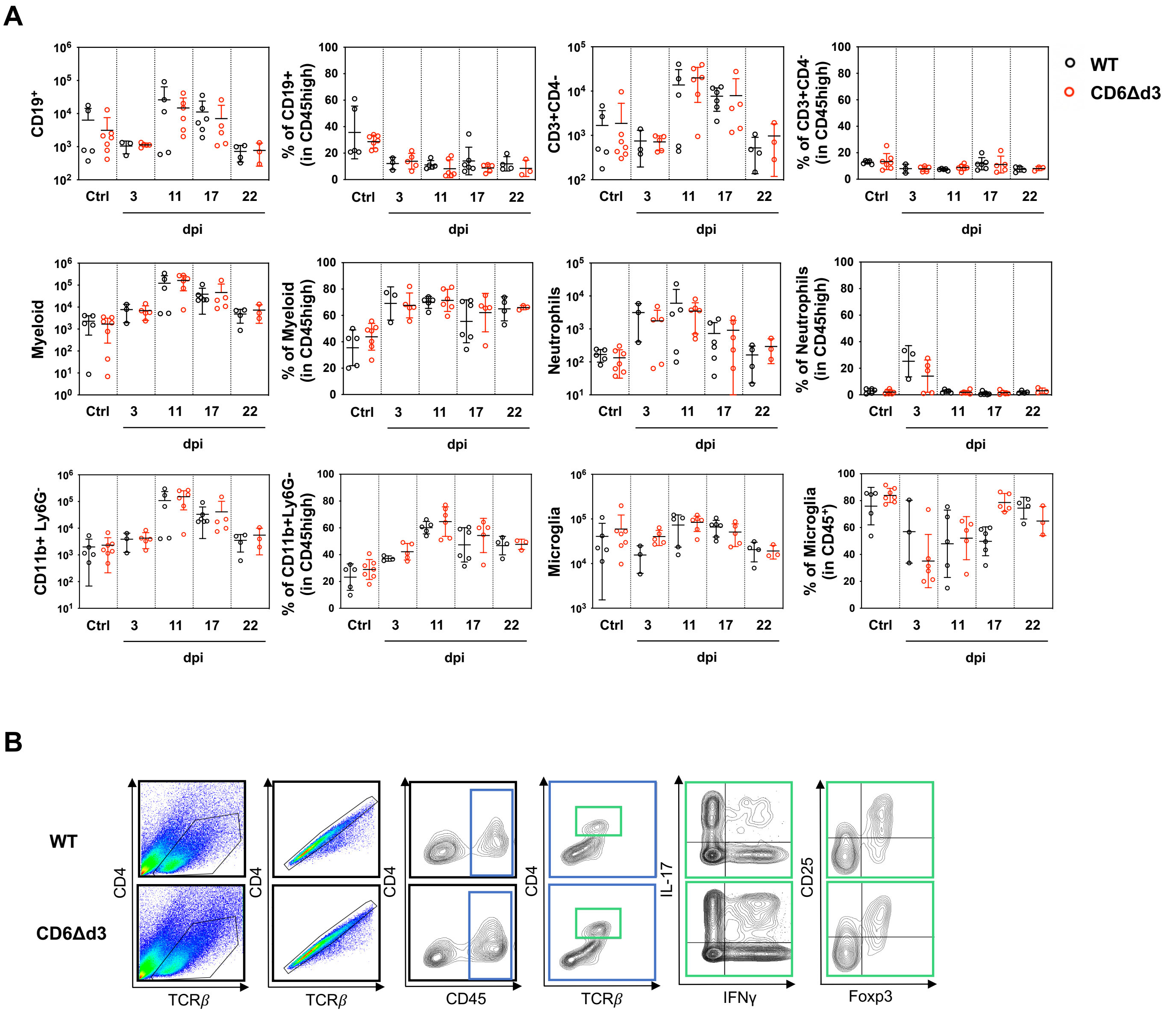


**Fig. S5. CNS cellular infiltration during the course of EAE.** (**A**) Flow cytometry analysis of total infiltrating cells in the brain parenchyma: CD19$^+$ (B cells); CD3$^+$CD4$^-$ (CD8 T cells); CD45$^{high}$CD11b$^+$ (all myeloid cells); CD45$^{high}$CD11b$^+$Ly6G$^+$ (neutrophils); CD45$^{high}$CD11b$^+$Ly6G$^-$ (macrophages, dendritic cells) and CD45$^{int}$CD11b$^+$ (microglia), at 0, 3, 11, 17 and 22 days post EAE induction. WT in black circles; CD6Δd3 in red circles. Ctrl: control (before induction - day zero); dpi: days post induction. (**B**) Representative flow cytometry plot scheme showing the gating strategy for CD4$^+$ T cell subsets in the brain parenchyma. FSC: forward scatter.
