## Supplementary Tables for "The dual character of the inhibitory functions of CD6"

**Movie S1. CD6 and CD3 co-localization at the immunological synapse.** Representative 3D projection of confocal microscopy z-stacks upon conjugate formation between sAg-loaded Raji without (A) or with CD166 (B and C) and E6.1 Jurkat cells expressing CD6WT (A and B) and CD6Δd3 (C). CD3 is shown in green and CD6 is displayed in red. Merge images allow to visualize CD3 and CD6 co-localization. 3D projection were done using ImageJ software and 3D Viewer plugin. Magnification: 63 x.

**Supplementary Table 1. Primers for CD6 constructs.** Underlined sequences correspond to restriction sites used for cloning.

| Construct | Primers (5'-3') Forward | Reverse |
|---|---|---|
| WT | ATAGGATCCATGTGGCTCTTCTTCGGGATC | ATAGCGGCCGCCTAGGCTGCGCTGATGTC |
| Δcyt | ATAGGATCCATGTGGCTCTTCTTCGGGATC | ATAGCGGCCGCCTATCCTTTAATTCTCAAGAG |
| Δd3 | ATAGGATCCATGTGGCTCTTCTTCGGGATC | CTGAGCACACCGCGCCCG |
| | ACGCGGGCGCGGTGTGCTCAGCTTCCCGGAGTTTGCACA | ATAGCGGCCGCCTAGGCTGCGCTGATGTC |
| Δd3Δcyt | ATAGGATCCATGTGGCTCTTCTTCGGGATC | CTGAGCACACCGCGCCCG |
| | ACGCGGGCGCGGTGTGCTCAGCTTCCCGGAGTTTGCACA | ATAGCGGCCGCCTATCCTTTAATTCTCAAGAG |
| D571stop | ATAGGATCCATGTGGCTCTTCTTCGGGATC | ATAGCGGCCGCCTACTCCCCTGAAGAGGTGCTCGA |
| D485stop | ATAGGATCCATGTGGCTCTTCTTCGGGATC | ATAGCGGCCGCCTATGAGTCCGAGCCAGAGTCTGA |
| Y430F | ATAGGATCCATGTGGCTCTTCTTCGGGATC | TACGGGGAGGGCAAATTTTCCTTTAAT |
| | ATTAAAGGAAAATTTGCCCTCCCCGTA | ATAGCGGCCGCCTAGGCTGCGCTGATGTC |
| Y452F | ATAGGATCCATGTGGCTCTTCTTCGGGATC | GGGGACCGGTTGAAAGCTATTGCTCCC |
| | GGGAGCAATAGCTTTCAACCGGTCCCC | ATAGCGGCCGCCTAGGCTGCGCTGATGTC |
| Y486F | ATAGGATCCATGTGGCTCTTCTTCGGGATC | GTCATAGTGCTCAAAGTCTGAGTCCGA |
| | TCGGACTCAGACTTTGAGCACTATGAC | ATAGCGGCCGCCTAGGCTGCGCTGATGTC |
| Y489F | ATAGGATCCATGTGGCTCTTCTTCGGGATC | GGCGCTGAAGTCAAAGTGCTCATAGT |
| | ACTATGAGCACTTTGACTTCAGCGCC | ATAGCGGCCGCCTAGGCTGCGCTGATGTC |
| Y503F | ATAGGATCCATGTGGCTCTTCTTCGGGATC | CCGCTGGGAATTGAAGAAGGTGGTCAG |
| | CTGACCACCTTCTTCAATTCCCAGCGG | ATAGCGGCCGCCTAGGCTGCGCTGATGTC |
| Y556F | ATAGGATCCATGTGGCTCTTCTTCGGGATC | GCTCCTCGGGTGAAACTGAGGGCCCAG |
| | CTGGGCCCTCAGTTTCACCCGAGGAGC | ATAGCGGCCGCCTAGGCTGCGCTGATGTC |
| Y572F | ATAGGATCCATGTGGCTCTTCTTCGGGATC | GGGACTATTGCAGAAATCCTCCCCTGA |
| | TCAGGGGAGGATTTCTGCAATAGTCCC | ATAGCGGCCGCCTAGGCTGCGCTGATGTC |
| Y629F | ATAGGATCCATGTGGCTCTTCTTCGGGATC | CTGGAAGTTCTGGAACCACTCCCCGGA |
| | CTGGAAGTTCTGGAACCACTCCCCGGA | ATAGCGGCCGCCTAGGCTGCGCTGATGTC |
| Y662F | ATAGGATCCATGTGGCTCTTCTTCGGGATC | ATAGCGGCCGCCTAGGCTGCGCTGATGTCATCGAAGTCATCGTTGTC |

**Table S2. Panel of antibodies for immunophenotyping of thymus and spleen.**

### Immunophenotyping Panels

| | Specificity | Clone | Provider |
|---|---|---|---|
| Thymus | CD003e | 2C11 | eBioscience |
| | CD004 | RM4.5 | BD Biosciences |
| | CD005 | 53-7.3 | BD Biosciences |
| | CD008a | 53-6.7 | BD Biosciences |
| | CD024 | M1.69 | BD Biosciences |
| | CD025 | PC61 | Biolegend |
| | CD027 | LG.3A10 | BD Biosciences |
| | CD044 | IM7 | Biolegend |
| | CD069 | H1.2F3 | BD Biosciences |
| | CD071 | R17217 | Biolegend |
| | CD117 | 2B8 | BD Biosciences |
| | CD161 | PK136 | BD Biosciences |
| | TCRd | GL3 | eBioscience |
| | QA-2 | 695H1.9.9 | Biolegend |
| Spleen | CD003e | 2C11 | BD Biosciences |
| | CD004 | RM4.5 | BD Biosciences |
| | CD005 | 53-7.3 | BD Biosciences |
| | CD008 | 53-6.7 | BD Biosciences |
| | CD011b | M1.70 | BD Biosciences |
| | CD011c | HL3 | BD Biosciences |
| | CD019 | 1D3 | BD Biosciences |
| | CD044 | IM7 | BD Biosciences |
| | CD062L | MEL14 | Biolegend |
| | CD161 | PK136 | BD Biosciences |
| | CD317 | 927 | eBioscience |
| | Ly6C | AL21 | BD Biosciences |
| | Ly6G | 1A8 | BD Biosciences |
| | TCRd | GL3 | eBioscience |
| | F4/80 | BM8 | Biolegend |
| | MHCII | M5-114.15.2 | Biolegend |
| | IgD | 11-26c.2A | Biolegend |

### Populations definition

| | | | | | | | | | | | | |
|---|---|---|---|---|---|---|---|---|---|---|---|---|
| Thymus | NKc | CD161+ | CD3e- | | | | | | | | | |
| | NK-Tc | CD161+ | CD3eint | | | | | | | | | |
| | DP cells | CD161- | CD27+ | CD4+ | CD8a+ | | | | | | | |
| | CD4 SP | CD161- | CD27+ | CD4+ | CD8a- | | | | | | | |
| | CD8 SP | CD161- | CD27+ | CD4- | CD8a+ | | | | | | | |
| | DN | CD161- | CD27+ | CD4- | CD8a- | | | | | | | |
| | TN | CD161- | CD27+ | CD4- | CD8a- | CD3e- | gd- | | | | | |
| | TN1 | CD161- | CD27+ | CD4- | CD8a- | CD3e- | gd- | CD44+ | CD25- | | | |
| | ETP (TN1a/b) | CD161- | CD27+ | CD4- | CD8a- | CD3e- | gd- | CD44+ | CD25- | CD24+ | CD117hi | |
| | TN1c | CD161- | CD27+ | CD4- | CD8a- | CD3e- | gd- | CD44+ | CD25- | CD24+ | CD117int | |
| | TN1d | CD161- | CD27+ | CD4- | CD8a- | CD3e- | gd- | CD44+ | CD25- | CD24+ | CD117- | |
| | TN1e | CD161- | CD27+ | CD4- | CD8a- | CD3e- | gd- | CD44+ | CD25- | CD24- | CD117- | |
| | TN2 | CD161- | CD27+ | CD4- | CD8a- | CD3e- | gd- | CD44+ | CD25+ | CD24+ | CD117+ | |
| | TN3a | CD161- | CD27+ | CD4- | CD8a- | CD3e- | gd- | CD44- | CD25+ | CD24+ | CD117- | CD71- |
| | TN3b | CD161- | CD27+ | CD4- | CD8a- | CD3e- | gd- | CD44- | CD25+ | CD24+ | CD117- | CD71+ |
| | TN4 | CD161- | CD27+ | CD4- | CD8a- | CD3e- | gd- | CD44- | CD25- | CD24+ | | |
| | immature DP | CD161- | CD27+ | CD4+ | CD8a+ | CD3e- | gd- | CD44- | CD25- | CD24+ | CD71+ | |
| | small resting DP | CD161- | CD27+ | CD4+ | CD8a+ | CD3e- | gd- | CD44- | CD25- | CD24+ | CD71- | |
| | CD3- CD69+ DP | CD161- | CD27+ | CD4+ | CD8a+ | CD3e- | gd- | CD44- | CD25- | CD24+ | CD71- | CD69+ |
| | CD3+ CD69+ DP | CD161- | CD27+ | CD4+ | CD8a+ | CD3e+ | gd- | CD44- | CD25- | CD24+ | CD71- | CD69+ |
| | CD3+ CD4 SP | CD161- | CD27+ | CD4+ | CD8a- | CD3e+ | gd- | CD44- | | | | |
| | Treg | CD161- | CD27+ | CD4+ | CD8a- | CD3e- | gd- | CD44- | CD25+ | | | |
| | Early CD4 SP | CD161- | CD27+ | CD4+ | CD8a- | CD3e- | gd- | CD44- | CD25- | CD24hi | | |
| | Late CD4 SP | CD161- | CD27+ | CD4+ | CD8a- | CD3e- | gd- | CD44- | CD25- | CD24lo | QA-2+ | |
| | CD3+ CD8 SP | CD161- | CD27+ | CD4- | CD8a+ | CD3e+ | gd- | CD44- | CD25- | | | |
| | Early CD8 SP | CD161- | CD27+ | CD4- | CD8a+ | CD3e+ | gd- | CD44- | CD25- | CD24hi | | |
| | Late CD8 SP | CD161- | CD27+ | CD4- | CD8a+ | CD3e+ | gd- | CD44- | CD25- | CD24lo | QA-2+ | |
| | gd-Tc | CD161- | CD27+ | CD4- | CD8a+ | CD3e+ | gd+ | | | | | |
| | DN3a gd-Tc | CD161- | CD27+ | CD4- | CD8a+ | CD3e+ | gd+ | CD25+ | CD71+ | | | |
| | DN4a gd-Tc | CD161- | CD27+ | CD4- | CD8a+ | CD3e+ | gd+ | CD25- | CD44- | CD71+ | | |
| | DN4b gd-Tc | CD161- | CD27+ | CD4- | CD8a+ | CD3e+ | gd+ | CD25- | CD44- | | | |
| | Cluster A gd Tc | CD161- | CD27+ | CD4- | CD8a+ | CD3e+ | gd+ | CD44+ | CD24- | | | |
| | Cluster B gd Tc | CD161- | CD27+ | CD4- | CD8a+ | CD3e+ | gd+ | CD44- | CD24+ | | | |
| Spleen | Neutrophils | CD11b+ | Ly6G+ | | | | | | | | | |
| | Eosinophils | CD11b+ | Ly6G- | Ly6Clo | SSChi | | | | | | | |
| | Macrophages | CD11bint | Ly6G- | Ly6C- | F4/80hi | CD11c- | | | | | | |
| | Monocytes | CD11b+ | Ly6G- | Ly6Chi | | | | | | | | |
| | pDC | CD11b- | Ly6G- | Ly6Chi | CD317+ | | | | | | | |
| | DC | CD11b- | Ly6G- | Ly6C- | CD317- | CD11c+ | MHCII+ | | | | | |
| | CD8a-Type DC | CD11b- | Ly6G- | CD11b+ | CD317- | CD11c+ | MHCII+ | CD8a+ | | | | |
| | CD11b-Type DC | CD11b- | Ly6G- | CD11b- | CD317- | CD11c+ | MHCII+ | CD11b+ | | | | |
| | NKc | CD161+ | Ly6G- | CD317- | CD5- | | | | | | | |
| | NK CD11b+Ly6C- | CD161+ | Ly6G- | CD317- | CD5- | CD11b+ | Ly6C- | | | | | |
| | NK CD11b+Ly6C+ | CD161+ | Ly6G- | CD317- | CD5- | CD11b+ | Ly6C+ | | | | | |
| | NK CD11b-Ly6C- | CD161+ | Ly6G- | CD317- | CD5- | CD11b- | Ly6C- | | | | | |
| | NK CD11b-Ly6C+ | CD161+ | Ly6G- | CD317- | CD5- | CD11b- | Ly6C+ | | | | | |
| | NK-Tc | CD161+ | Ly6G- | CD317- | CD5+ | | | | | | | |
| | Bc | CD161- | Ly6G- | CD317- | CD19+ | MHCII+ | | | | | | |
| | B1 Bc | CD161- | Ly6G- | CD317- | CD19+ | MHCII+ | CD5+ | | | | | |
| | B2 Bc | CD161- | Ly6G- | CD317- | CD19+ | MHCII+ | CD5- | | | | | |
| | IgDlo B2 Bc | CD161- | Ly6G- | CD317- | CD19+ | MHCII+ | CD5- | IgDlo | | | | |
| | IgD+ B2 Bc | CD161- | Ly6G- | CD317- | CD19+ | MHCII+ | CD5- | IgDhi | | | | |
| | Tc | CD161- | Ly6G- | CD317- | CD5+ | CD3+ | | | | | | |
| | gd-Tc | CD161- | Ly6G- | CD317- | CD5+ | CD3+ | TCRd+ | | | | | |
| | EM gd-Tc | CD161- | Ly6G- | CD317- | CD5+ | CD3+ | TCRd+ | CD44+ | CD62L- | | | |
| | Naive gd-Tc | CD161- | Ly6G- | CD317- | CD5+ | CD3+ | TCRd+ | CD44- | CD62L+ | | | |
| | CM gd-Tc | CD161- | Ly6G- | CD317- | CD5+ | CD3+ | TCRd+ | CD44+ | CD62L+ | | | |
| | ab Tc | CD161- | Ly6G- | CD317- | CD5+ | CD3+ | TCRd- | | | | | |
| | CD4 Tc | CD161- | Ly6G- | CD317- | CD5+ | CD3+ | TCRd- | CD4+ | | | | |
| | EM CD4 Tc | CD161- | Ly6G- | CD317- | CD5+ | CD3+ | TCRd- | CD4+ | CD44+ | CD62L- | | |
| | CM/Naive CD4 Tc | CD161- | Ly6G- | CD317- | CD5+ | CD3+ | TCRd- | CD4+ | CD44- | CD62L+ | | |
| | CD8 Tc | CD161- | Ly6G- | CD317- | CD5+ | CD3+ | TCRd- | CD8+ | | | | |
| | Naive CD8 Tc | CD161- | Ly6G- | CD317- | CD5+ | CD3+ | TCRd- | CD8+ | CD44- | CD62L+ | | |
| | CM CD8 Tc | CD161- | Ly6G- | CD317- | CD5+ | CD3+ | TCRd- | CD8+ | CD44+ | CD62L+ | | |
| | EM CD8 Tc | CD161- | Ly6G- | CD317- | CD5+ | CD3+ | TCRd- | CD8+ | CD44+ | CD62L- | | |
